# High-resolution Structures of multiple 5-HT_3A_R-setron complexes reveal a novel mechanism of competitive inhibition

**DOI:** 10.1101/2020.03.30.016154

**Authors:** Sandip Basak, Arvind Kumar, Steven Ramsey, Eric Gibbs, Abhijeet Kapoor, Marta Filizola, Sudha Chakrapani

## Abstract

Serotonin receptors (5-HT_3A_R) play a crucial role in regulating gut movement, and are the principal target of setrons, a class of high-affinity competitive antagonists, used in the management of nausea and vomiting associated with radiation and chemotherapies. Structural insights into setron-binding poses and their inhibitory mechanisms are just beginning to emerge. Here, we present high-resolution cryo-EM structures of full-length 5-HT_3A_R in complex with palonosetron, ondansetron, and alosetron. Each structure reveals a distinct interaction fingerprint between the setron and binding-pocket residues that may underlie their diverse affinities. In addition, setrons elicit varying degrees of conformational change throughout the channel that, quite surprisingly, lie along the channel activation pathway, suggesting a novel mechanism of competitive inhibition. Molecular dynamic simulations were used to assess binding-poses and the drug-target interaction dynamics. Together, this study provides a molecular basis for setron binding affinities and their inhibitory effects.

## Introduction

Cancer treatments by radiation or chemotherapy triggers the release of excess serotonin from the mucosal enterochromaffin cells in the upper gastrointestinal tract^1^. Serotonin binds to serotonin (3) receptors (5-HT_3_Rs), a pentameric ligand-gated ion channel (pLGIC), on the vagal afferent nerve in the gut and on the chemoreceptor trigger zone in the brainstem leading to severe nausea and vomiting in the patients. These common side effects of cancer treatment take a significant physical and psychological toll on cancer patients. Without management, these side effects can reduce patient compliance, undermining treatment success^2^. Furthermore, uncontrolled debilitating side-effects result in secondary complications such as dehydration and anorexia that require additional hospitalization and increase overall healthcare costs.

Current antiemetic therapies include a 5-HT_3_R antagonist treatment regimen, which is considered a major advancement in improving patient quality of life during cancer treatment. Setrons, competitive antagonists of 5-HT_3_R, are effective in the prevention of chemotherapy-induced nausea and vomiting (CINV), radiation therapy-induced nausea and vomiting (RINV), and postoperative nausea and vomiting (PONV) ^3,4^. CINV nausea occurs in acute and delayed phases. The first generation of FDA approved setrons are effective for treating acute but not delayed phase nausea due to their short plasma half-lives. They belong to the following major classes based on their chemical structures: carbazole (*e.g.* ondansetron), indazole (*e.g.* granisetron), indole (*e.g.* dolasetron, tropisetron), and imidazole (*e.g.* alosetron). Although setrons share the same fundamental mechanism of action, they have varying efficacies, dose-response profiles, duration of action, and off-target responses. These differences perhaps underlie variable patient response, particularly in the context of acute and refractory emesis^5^. Palonosetron, the only FDA approved second generation setron, a derivative of isoquinoline, is shown to have a longer half-life, improved bioavailability, and efficacy. In addition, palonosetron is implicated in causing receptor internalization, which further improves anti-emetic properties. Beyond their role in controlling emesis, setrons are used to treat GI disorders including irritable bowel syndrome (IBS), obesity, and several inflammatory, neurological and psychiatric disorders such as migraine, drug abuse, schizophrenia, depression, anxiety, and cognitive disorders. However, in some cases, toxicity and adverse side-effects have hampered their use. For example, with alosetron, which was approved by the FDA for the treatment of diarrhea-predominant IBS, led to severe ischemic colitis in many patients^6^. Given the broad therapeutic potential of 5-HT_3A_R antagonists, the prospect of substantial therapeutic gains by probing the setron pharmacophore is encouraging, as well as developing novel pharmaceuticals with higher efficacy and reduced side-effects.

At the physiological level, the 5-HT_3_Rs play an important role in gut motility, visceral sensation, and secretion ^7-12^, and are also implicated in pain perception, mood, and appetite. 5-HT_3_Rs are the only ion channels^13^ among the large family of serotonin receptors, the rest being GPCRs. 5-HT_3_Rs are expressed as homopentamers of subunit A or heteropentamers of subunit A, in combination with B, C, D, or E subunits^14^. Compositional and stoichiometric differences lead to differential responses to serotonin, gating kinetics, permeability, and pharmacology^15-17^. This functional diversity, tissue specific expression patterns, and distinct pathophysiology of 5-HT_3_R isoforms establish a need for subtype specific drugs to address diverse clinical needs^18^. Of note, granisetron, ondansetron, palonosetron, and alosetron have slightly different affinities for various receptor subtypes^19^. Ondansetron, in addition to binding to 5-HT_3_Rs, also binds to several GPCRs, such as 5-HT_1B_R, 5-HT_1C_R, α1-adrenergic receptors and μ-opioid receptors^20^. Granisetron binds to all subtypes of 5-HT_3_R, but has little or no affinity for 5-HT_1_R, 5-HT_2_R and 5-HT_4_R receptors. Palonosetron is highly selective for 5-HT_3A_R and dolasetron for 5-HT_3B_R^21^. Structural insights into setron-binding poses came initially from crystal structures of the acetylcholine binding protein (AChBP), bound to granisetron, tropisetron, or palonosetron^22-24^ and more recently from 5-HT_3A_R complexed with tropisetron and granisetron^25,26^. While some of the basic principles of setron-binding are now clear, there is still limited understanding of differing pharmacodynamics among setrons and the associated clinical relevance.

In the present study, we have solved cryo-EM structures of the full-length 5-HT_3A_R in complex with palonosetron, ondansetron, and alosetron at the resolution range of 2.9 Å to 3.3 Å. Together with our previously solved structure of 5-HT_3A_R in complex with granisetron (grani-5-HT_3A_R), we provide details of various setron-binding modes and the ensuing conformational changes that lead to channel inhibition. Using molecular dynamics (MD) simulations and electrophysiology, we have further validated setron-binding modes and interactions within the conserved binding pocket. Combined with abundant functional, biochemical, and clinical data, these new findings may serve as a structural blueprint of drug-target interactions that can guide new drug development.

## Results

### Cryo-EM structures of setron-5-HT_3A_R complexes

Structures of the full-length 5-HT_3A_R in complex with setrons were solved by single-particle cryo-EM. Detergent solubilized 5-HT_3A_R was incubated with 100 µM of palonosetron, ondansetron, or alosetron for 1 hour prior to vitrification on cryo-EM grids. Iterative classifications and refinement produced a final three-dimensional reconstruction at a nominal resolution of 3.3 Å for 5-HT_3A_R-Palono (with 91,163 particles), 3.0 Å for 5-HT_3A_R-Ondan (67,333 particles), and 2.9 Å for 5-HT_3A_R-Alo (46,065 particles) (Supplemental Figures 1a and 1b). The local resolution of the map was estimated using ResMap and in the range of 2.5-3.5 Å for each of these reconstructions (Supplemental Figure 1c). Structural models were built using refined maps containing density for the entire extracellular domain (ECD), transmembrane domain (TMD), and the structured regions of the intracellular domain (ICD) (Fig. 1a and Supplemental Figure 2). Overall, each of setron-bound 5-HT_3A_R complexes has an architecture similar to previously solved 5-HT_3A_ receptors^25,27-29^. Among the non-protein densities present in the map at the ECD are the three sets of peripheral protrusions corresponding to N-linked glycans (Fig. 1a, right) and a strong, unambiguous density at each of the intersubunit interfaces, corresponding to individual setrons (Figs. 1b). Besides this site, no additional densities for setrons were found under these conditions, although there have been predictions that palonosetron may act as both an orthosteric and allosteric ligand^30^.

**Figure 1.**
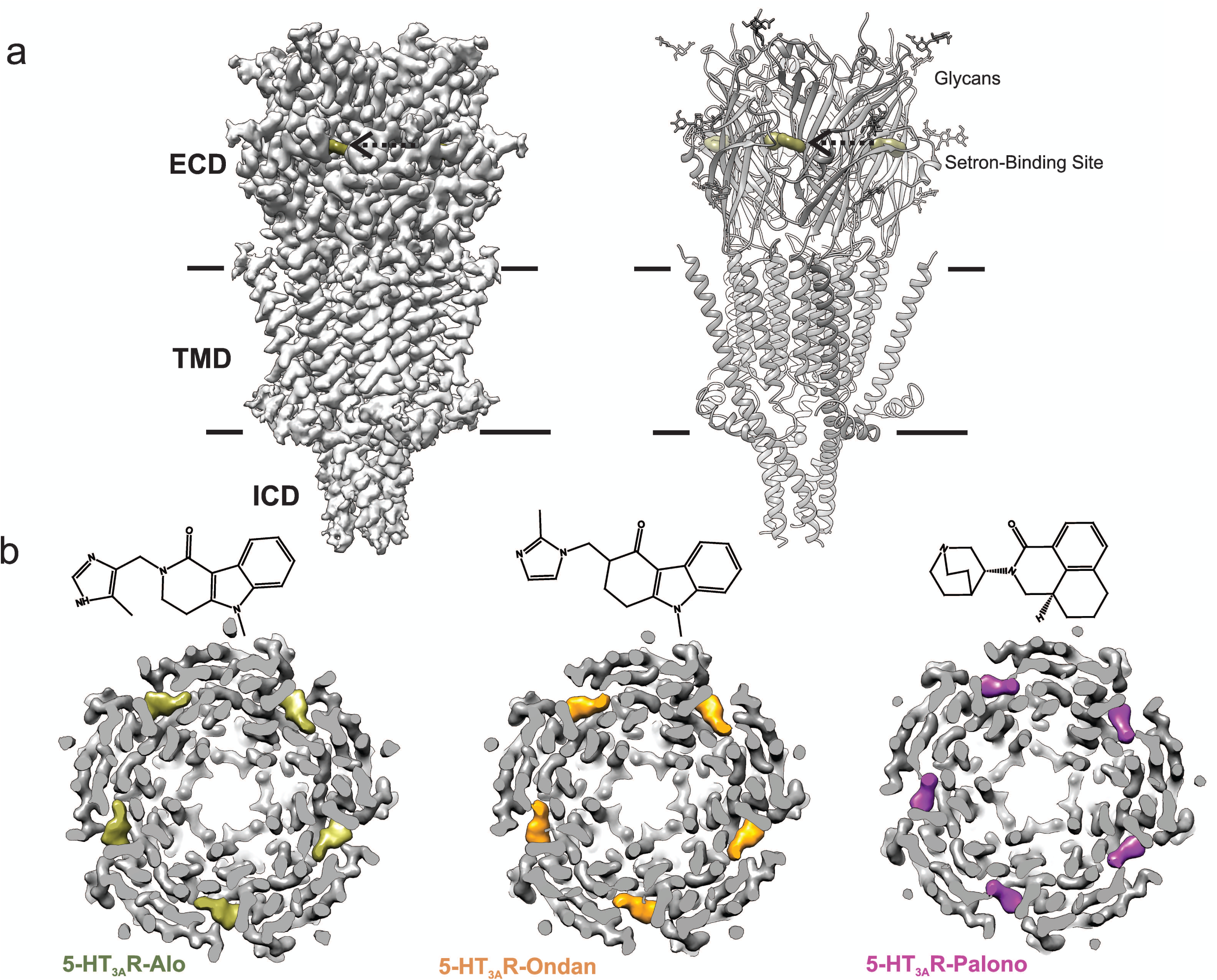
Cryo-EM structure of 5-HT_3A_R-setron complexes. **a** Three dimensional reconstruction of 5-HT_3A_R-Alo at 2.92 Å resolution (*left*) and the corresponding structural model (*right*) that shows the overall architecture consisting of the extracellular domain (ECD), transmembrane domain (TMD), and structural regions of the intracellular domain (ICD). The alosetron density is shown in *deep olive* color and the three sets of glycans are shown as stick representation. Arrow points towards the setron density. Solid line denote putative membrane limits. **b** Extracellular view of 5-HT_3A_R-Alo (*left*), 5-HT_3A_R-Ondan (*middle*), and 5-HT_3A_R-Palono (*right*) maps sliced at the neurotransmitter-binding site. In each case, the five molecules of respective setrons are highlighted in colors. Chemical structures of setrons are shown above.

The map quality was particularly good at the ligand-binding site allowing us to model sidechains and the setron orientation. Setrons bind within the canonical neurotransmitter binding pocket and are lined by residues from Loops A, B, and C on the principal (+) subunit and Loops D, E, and F from the complementary (-) subunit (Fig. 2). Residues within 4 Å of setron include Asn101 in Loop A, Trp156 in Loop B, Phe199 and Tyr207 in Loop C, Trp63 and Arg65 in Loop D, and Tyr126 in Loop E. These residues are strictly conserved, and perturbations at each of these positions impact efficacy of setrons and serotonin^31-33^. In each setron-5-HT_3A_R complex, the essential pharmacophore of setron is placed in a similar orientation: the basic amine is at the deep-end of the pocket in the principal subunit; the defining aromatic moiety interacts with residues in the complementary subunit; and the carbonyl-based linker, between the two groups, is essentially coplanar with the aromatic ring.

**Figure 2.**
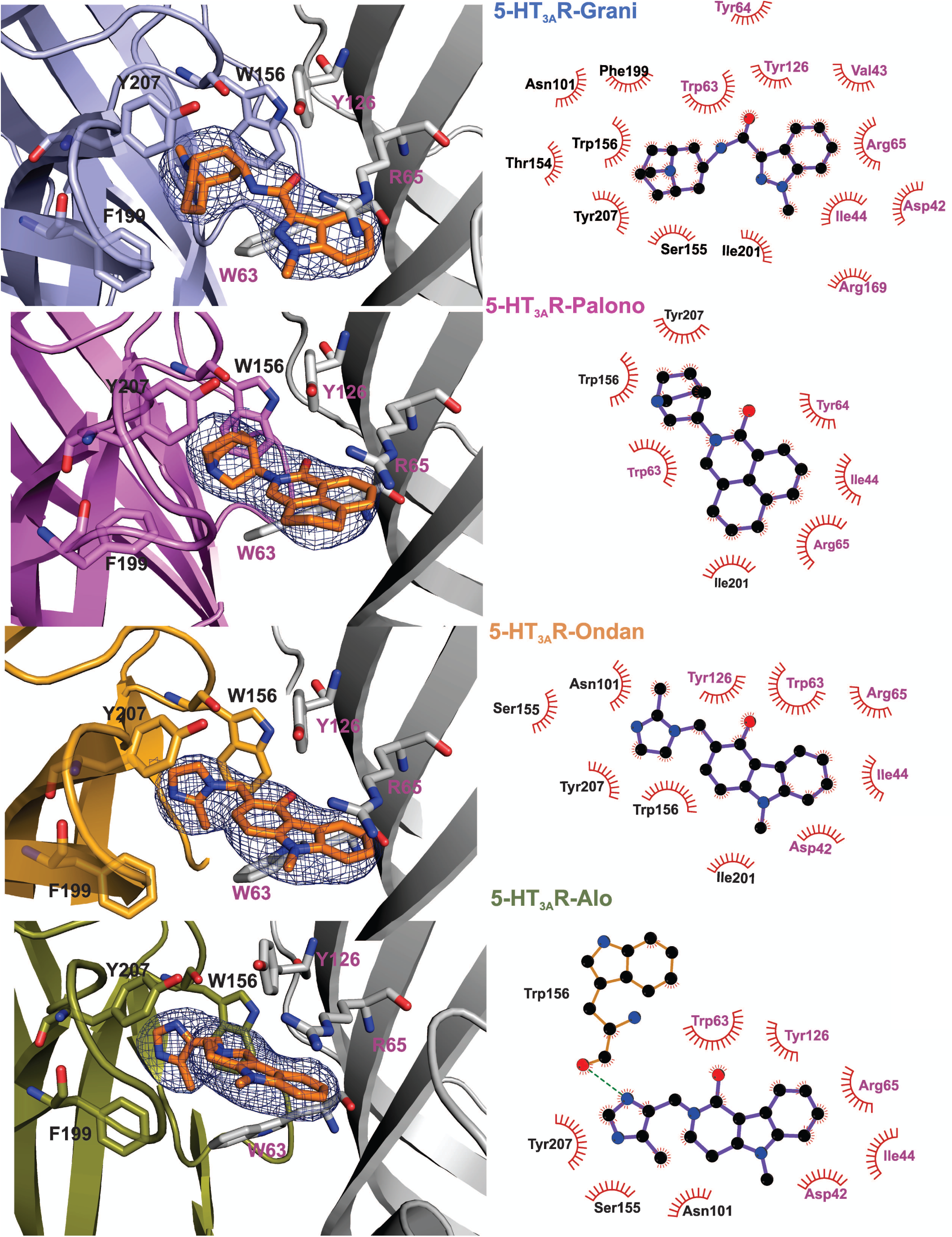
Setron-binding poses. **a** Cryo-EM density for the setrons, located at the canonical neurotransmitter-binding site. The map is contoured at 9*σ* (5-HT_3A_R-Grani)^26^; 8.5*σ* (5-HT_3A_R-Palono); 7*σ* (5-HT_3A_R-Ondan); 6*σ* (5-HT_3A_R-Alo). The binding-site lies at the interface of the principal (colored) and the complementary (gray) subunits. The binding site residues are shown in stick representation with residues from the principal subunit labeled in *black* and those from the complementary subunit in *magenta*. From *top to bottom*: 5-HT_3A_R-Grani, 5-HT_3A_R-Palono, 5-HT_3A_R-Ondan, and 5-HT_3A_R-Alo. **b** LigPlot analysis of setron-5-HT_3A_R interactions. Most interactions are hydrophobic in nature. Putative hydrogen bond between Trp156 and alosetron is shown as a *green* dotted line.

The basic amine of the setron is in a bicyclic ring in granisetron and palonosetron, and a diazole ring in ondansetron and alosetron. The amine is within 4 Å of Trp156 (loop B), Tyr207 (loop C), Trp63 (loop D) and Tyr126 (loop E), and are likely to be involved in polar interactions with these residues. In particular, the carbonyl oxygen of Trp156 is close to the amine group of setron, and in the 5-HT_3A_R-Alo, it forms a hydrogen bond with the amine group in the diazole ring. The relative orientation of the tertiary nitrogen and Trp156 is conducive for a cation-pi interaction, as seen in the AChBP-5-HT_3_ chimera structure^22^. A similar interaction is also predicted for the primary amine group of serotonin^34^. The aromatic, hydrophobic end of the molecule is an indazole in granisetron, isoquinoline in palonosetron, carbazole in ondansetron, and imidazole in alosetron. It is oriented toward the complementary subunit, and lies parallel to the membrane. In this orientation, the aromatic moiety is stabilized by a number of hydrophobic interactions with Ile44, Trp63, Tyr64, Ile201, and Tyr126. The setron molecule is within 4-5 Å and potentially makes π-π interactions (edge-to-face or face-to-face) with Trp63, Tyr126, Trp156, and Tyr207. These interactions are also consistent with our MD simulations (discussed below). In addition, the planar aromatic rings lie beneath Arg65, and in close proximity to the positively charged nitrogen in the guanidinium group of Arg65, revealing a potential cation-pi interaction (Fig. 3a). This interaction was also observed in the AChBP-5-HT_3_ chimera^22^ and 5-HT_3A_R-Grani structures^26^. As previously noted in 5-HT_3A_R-Grani, the setron position causes reorientation of Arg65 (β2 strand or loop D) and Trp168 (β8-β9; loop F)^26^. Earlier reports also predicted large orientational differences for Trp168 when the binding-site was occupied by agonist or antagonist^35^. In this position, Arg65 is in a network of interactions involving Asp42 (β1), Try126 (β6), Trp168 (β8-β9; loop F), Arg169 (β8-β9; loop F), and Asp177 (β8-β9; loop F) (Fig. 3b). Glu102 (loop A) which is in the vicinity of ligand binding site is in a hydrogen bond network with Thr133 and Ala134 carbonyl (β6 strand). Interestingly, both of these networks are also present in serotonin-bound 5-HT_3A_R, but absent in 5-HT_3A_R-Apo, indicating the ligand-induced formation of the interaction network^28,29^.

**Figure 3.**
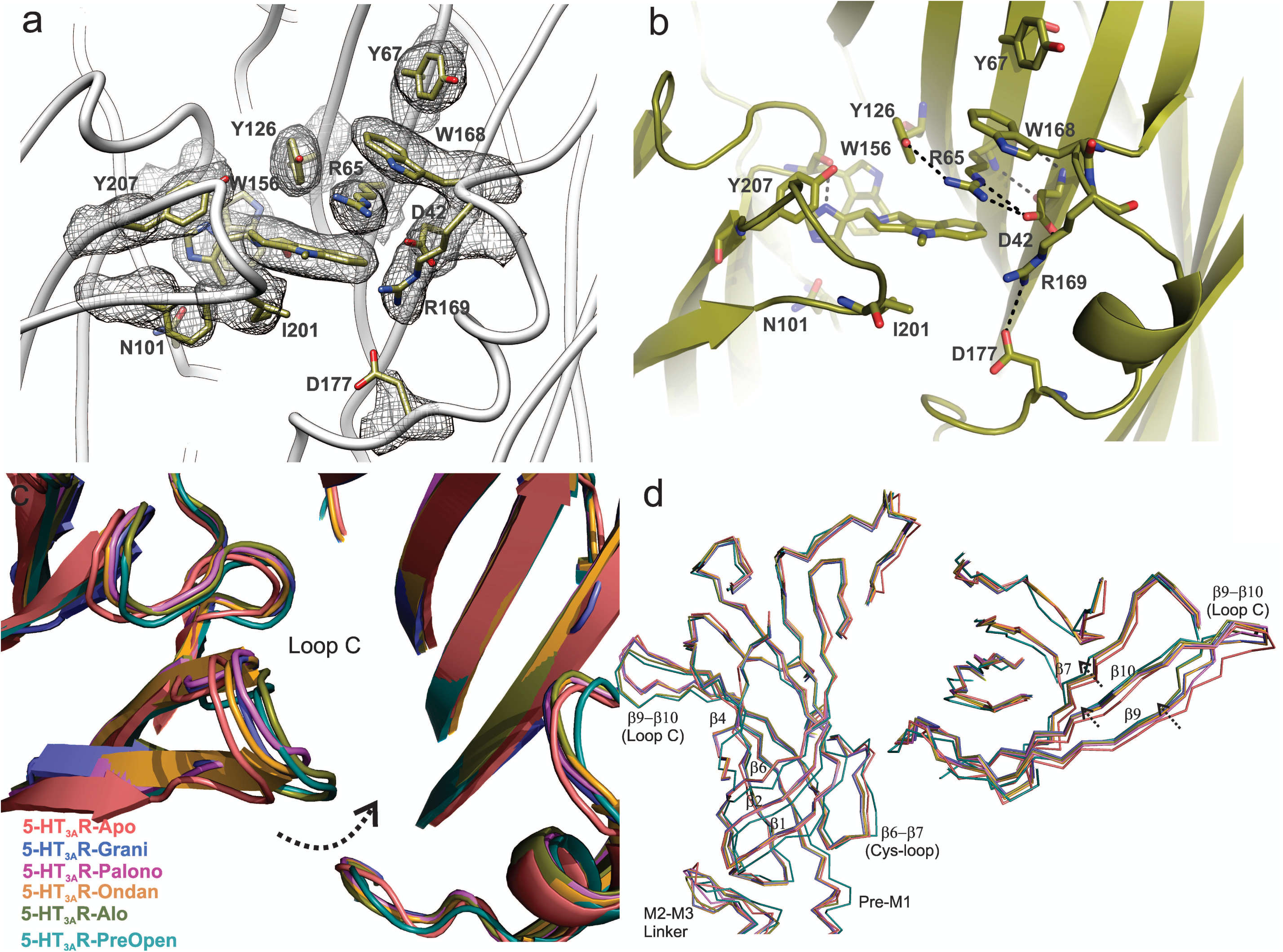
Setron-binding pocket and conformational changes in Loop C. **a** Arrangement of residues lining the binding site as seen in 5-HT_3A_R-Alo. The density map around the binding-site residues are contoured at 9*σ.* **b** A *close-up, side-view* of the subunit-interface reveals extensive interactions between residues on the complementary subunit. Side-chains which are within 4 Å distance are marked by dotted line. **c** Global alignment of 5-HT_3A_R-Apo, 5-HT_3A_R-State1 (serotonin-bound), 5-HT_3A_R-Grani, 5-HT_3A_R-Palono, 5-HT_3A_R-Ondan, and 5-HT_3A_R-Alo structures. With respect to 5-HT_3A_R-Apo, the serotonin- and setron-bound conformations reveal an inward positioning of Loop C (*shown by arrow*). **d** Relative displacement of the inner β-strands seen from a side-view (*left panel*) and the outer β-strands seen from the top (*right panel*). Arrows indicate the direction of movement.

Our initial expectation was that, as a highly-potent competitive antagonist, setrons would stabilize a 5-HT_3A_R-Apo like conformation. However, 5-HT_3A_R-Grani revealed a counter-clockwise twist of beta strands in the ECD leading to a small inward movement of Loop C (connecting β9-β10 strands) closing-in on granisetron. The Loop C conformation has been correlated to the nature of the ligand in the binding site and agonist efficacy. The AChBP-ligand complexes have shown that agonist binding induces a “closure” of Loop C, capping the ligand-binding site^36^. This conformational change may be part of a conserved pLGIC mechanism that couples ligand binding to channel opening through the ECD-TMD interfacial loops. Antagonist-bound structures show Loop C further extended outward^36^, while partial agonists seem to induce partial Loop C closure but not to the level achieved by agonists^23^. The 5-HT_3A_R and other pLGIC structures solved thus far, in the apo and agonist-bound states, follow this general trend ^25,28,29^. However, studies have shown that unliganded pLGIC gating kinetics remain unaffected by Loop C truncation^37^, raising ambiguity over its role in the channel opening mechanism. In comparison to the 5-HT_3A_R-Apo, the Loop C conformation in the 5-HT_3A_R-setron structures are positioned inward to a varying degree, and in the 5-HT_3A_R-Alo the orientation is similar to the serotonin-bound conformation (State-I)^26^ (Fig. 3c). The twisting inward movement does not pertain to Loop C alone, it was also seen in adjoining β7, β9 and β10 strands that form the outer-sheets of the β-sandwich core, with notable deviation from 5-HT_3A_R-Apo in the vicinity of the binding pocket (Fig. 3d, *right panel*). There are minimal changes in the β-strands of the inner sheets (β1, β2, β6) (Fig. 3d, *left panel;* Supplemental Figure 3). These conformational changes approach those seen in serotonin bound 5-HT_3A_R (State-I)^29^.

Conformational changes are also present in the TMD and may arise from small twisting movements in the ECD. Interestingly, in each of the 5-HT_3A_R-setrons structures, the pore-lining M2 helices are positioned away from the central axis, and are in a more-expanded conformation than the 5-HT_3A_R-Apo (Fig. 4a). At positions Val260 (13′), Leu260 (9′), Ser 253 (2′), and Glu250 (−1′), where the pore is constricted to below the hydrated Na^+^ radii^38^ in 5-HT_3A_R-Apo, the pore-radii is larger in 5-HT_3A_R-setron structures (Fig. 4b). However, these conformations are expected to remain non-conducting because Leu260 (9′) is constricted to ∼2.3 Å. While there are small conformational changes in the ICD, the post-M3 loop occludes the lateral portals at the interface of the TMD and ICD which are ion exit paths. The extent of occlusion is similar to what was seen in 5-HT_3A_R-apo^28^.

**Figure 4.**
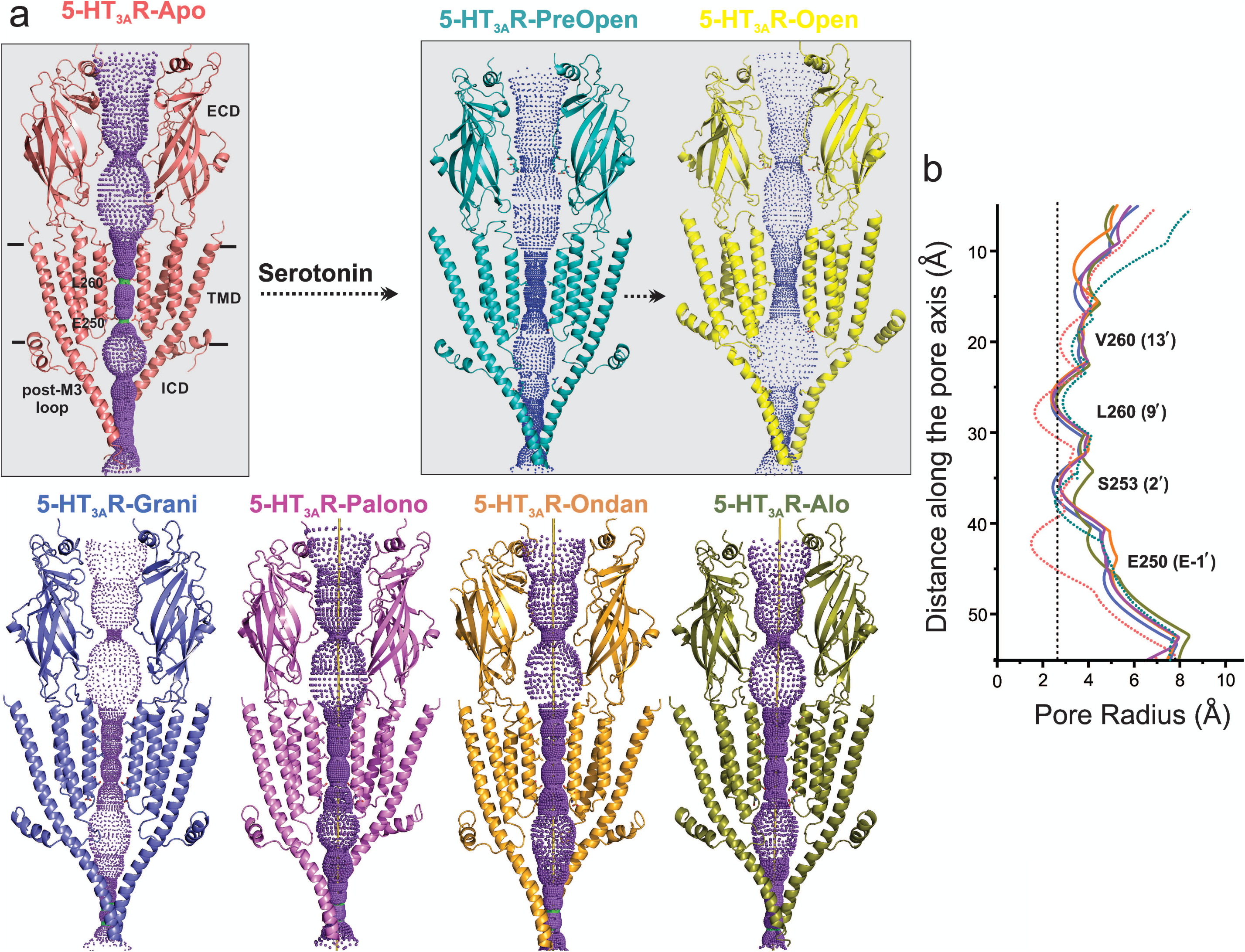
Pore profiles of 5-HT_3A_R in Apo, serotonin-, and setron-bound states. **a** Ion conduction pathway predicted by HOLE^52^. Models are shown in cartoon representation. Only two subunits are shown for clarity. The locations of pore constrictions are shown as sticks. **b** The pore radius is plotted as a function of distance along the pore axis. The dotted line indicates the approximate radius of a hydrated Na^+^ ion which is estimated at 2.76 Å (right)^38^.

### Molecular Dynamics Simulations

Several structural analyses of 5-HT_3A_R-palono, 5-HT_3A_R-ondan, and 5-HT_3A_R-alo were performed by investigating 100 ns MD simulations of these complexes embedded in a 1-palmitoyl-2-oleoyl phosphatidyl choline (POPC) membrane and encased in water with 150 mM NaCl. To assess the stability of each setron binding-pose modeled from cryo-EM density, we quantified the root mean square deviation (RMSD) of each pose relative to its starting conformation, averaged across each subunit. We found that all setrons maintained a low RMSD (<2.5 Å), with palonosetron demonstrating the largest RMSD among the group (Fig 5a). During the simulation, the bicylic ring displayed considerable fluctuation and positional reorientation. In these positions, palonosetron had distinct interactions with binding site residues (Supplemental Figure 4). To evaluate the types of interactions that these setrons maintained with protein sidechains during MD simulation, we calculated 5-HT_3A_R-setron interaction fingerprints (see figure legend or methods for full interaction type definitions) for each protomer in the complex (Supplemental Figures 5 and 6). Hydrophobic interactions comprise the majority of these interaction fingerprints, with some contributions from water-mediated interactions and aromatic interactions. Notably, the alosetron fingerprints suggests that this compound forms stronger interactions with Asp202 and Trp156 when compared to palonosetron, ondansetron, and granisetron.

**Figure 5.**
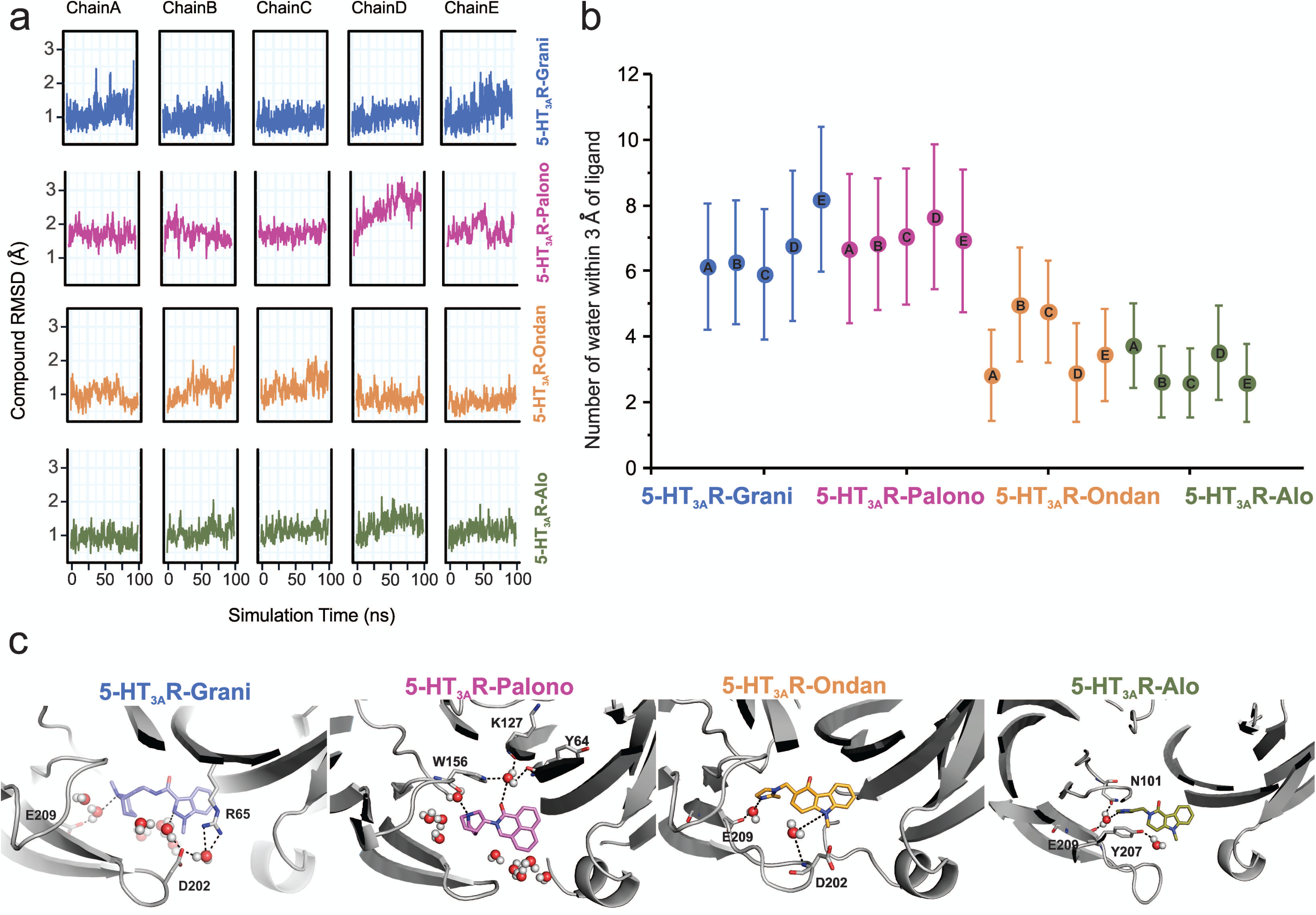
Assessment of conformation stability of setron-binding poses by molecular dynamic simulations. **a** Time evolution of Root Mean Square Deviation (RMSD) of setrons’ heavy atoms relative to their initial cryo-EM conformations of 5-HT_3A_R for each protomer subunit. **b** Assessment of the number of water molecules present within each setron-binding site during MD simulation. Shown are of the average number of water molecules (defined as a count of water oxygen atoms within 3 Å of any setron atoms) for each setron-bound simulation subdivided by protomer and the corresponding standard deviation. **c** Snapshots during the simulation showing water molecules in the pocket.

To characterize the flexibility of Loop C in our 5-HT_3A_R-setron simulations, we evaluated a number of structural quantities. First, we assessed the RMSD of Loop C for each protomer in each 5-HT_3A_R-setron simulation by evaluating the distances of Cα, carbonyl carbon, and backbone nitrogen atoms of residues Ser200 through Asn205 referenced to their initial cryo-EM conformations (Supplemental Figure 7a). These data suggest that Loop C is stable in its initial cryo-EM conformation, particularly in the 5-HT_3A_R-Alo and 5-HT_3A_R-Ondan simulations, but can also adopt an alternate ‘open’ conformation where Loop C extends away from the binding site surface. To further evaluate Loop C conformational flexibility in our MD simulations we defined a custom dihedral formed by the Cα atoms of residues Ala208, Phe199, Glu198, and Ile203 that measured the orientation of the loop relative to the principal protomer binding site (Supplemental Figure 7b). This dihedral was defined in such a way that a large angle would denote that the loop is oriented away from the binding site and small or negative angles indicate that the loop is oriented towards the binding site. This data demonstrates that Loop C tends to remain in its initial cryo-EM resolved conformation, particularly in the 5-HT_3A_R-Alo MD simulation.

To assess the impact of Loop C movement on the relative size of each setron-binding pocket, we quantified the number of water molecules found within each binding site. This was evaluated by counting water oxygen atoms within 3 Å of any setron atom. Since each setron remained stable within its respective binding site, this measurement represents an approximation of binding-site volume. This data shows that alosetron and ondansetron have a lower number of water molecules within their binding sites (Fig 5b and 5c).

To understand the motion of Loop C between the ‘closed’ cryo-EM structure and the MD sampled ‘open’ conformation we evaluated the minimum polar side chain atom distance between Arg65 and Asp202, residues known to form a hydrogen bond interaction that may effectively rigidify loop C in a ‘closed’ conformation^39^. We hypothesized that in our MD simulations Loop C would not adopt an ‘open’ conformation if an Arg65-Asp202 interaction was formed. We find that the 5-HT_3A_R-Alo MD simulation maintained an interaction between Arg65 and Asp202 more often than in any other setron-bound structure, and that most setron-bound simulations did not appreciably form this stabilizing interaction (Fig 6a and 6b). Thus, our mechanistic hypothesis is such that when Arg65 is interacting with Asp202, Loop C is in a stable ‘closed’ conformation, which in turn reduces the accessibility of the binding pocket to water, and incidentally contributes to the higher stability of the ligand binding pose.

**Figure 6.**
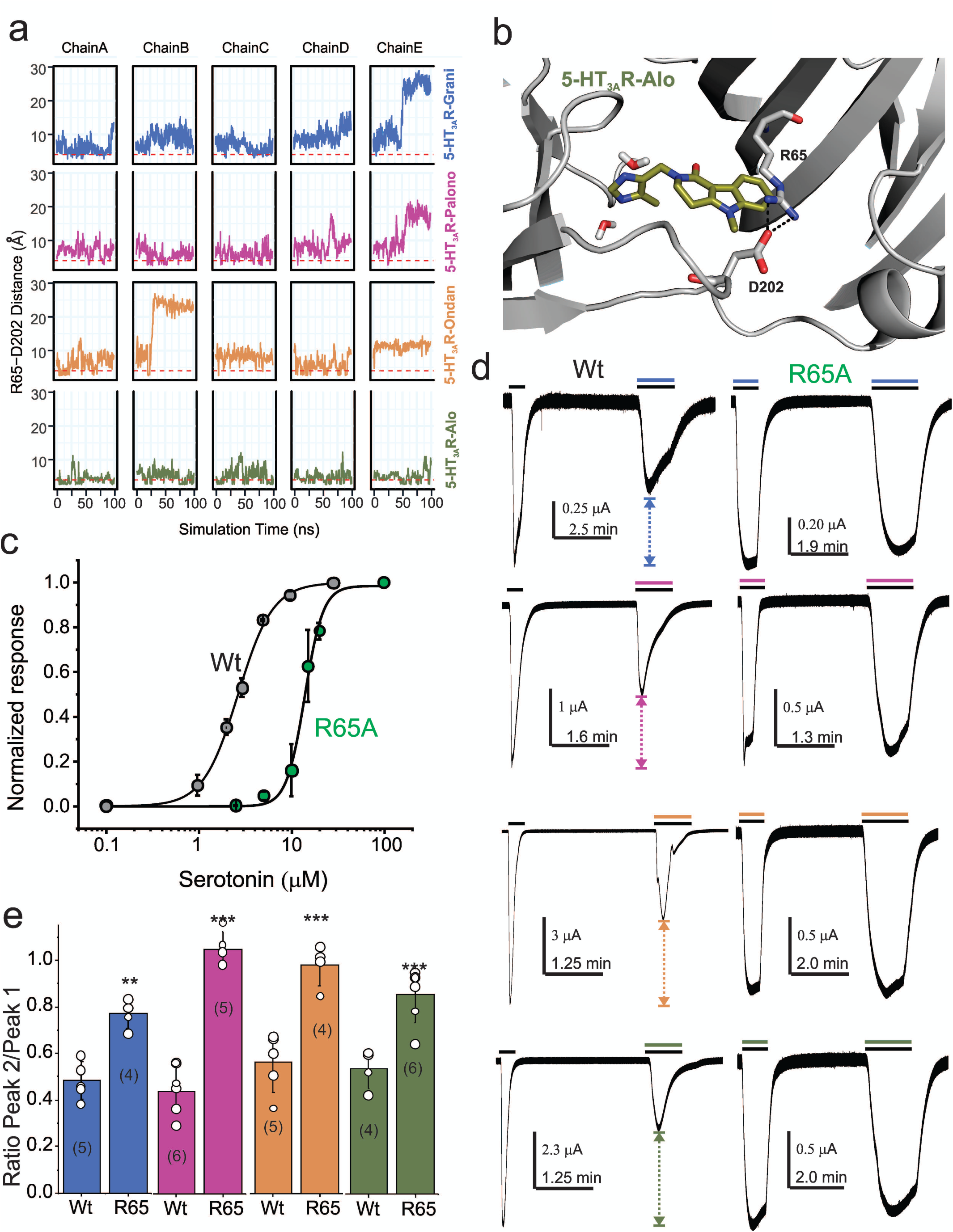
Dynamic interaction between Arg65 and Asp202. **a** Time evolution of the minimum distance between side chain polar atoms of Arg65 and Asp202 throughout 100 ns simulations. A 4 Å distance threshold is shown as a red dashed line to denote a generous cutoff for H-bond interactions between these residues. **b** MD snapshots that show the Arg65-Asp202 interaction. **c** Dose response curve for serotonin activation measured by TEVC recordings (at −60mV) for WT 5-HT_3A_R and R65A expressed in oocytes. The EC_50_, the Hill coefficient (nH), and the number of independent oocyte experiments are: WT (EC_50_: 2.70 ± 0.09 μM; nH: 2.3 ± 0.17; n: 3) and R65A (EC_50_: 13.79 ± 0.50 μM; nH: 4.4 ± 0.59; n: 4)^26^ **d** Functional analysis of perturbation of Arg65-Asp02 interaction. Currents were elicited in response to serotonin (concentrations used near EC_50_ values WT-2 μM, and R65A-10 μM) with and without co-application of setrons. Dotted arrows show the extent of setron inhibition in each case. **e** A plot of the ratio of peak current in the presence of setron to peak current in the absence of setron is shown for WT and R65A. Data are shown as mean ± s.d (n is indicated in parenthesis). Significance at p=0.01 (***) and p=0.05 (**) calculated by two sample t-test for wild type and R65A.

Our MD simulations predict that Arg65 may have a differential effect on the binding of various setrons. In agreement, mutations at the Arg65 position in human 5-HT_3A_R abolish granisetron binding but tropisetron binding is only reduced^40^. To further assess the role of Arg65 in binding various setrons, we measured the extent of inhibition of serotonin-induced currents. Since for competitive antagonists, the extent of inhibition depends on agonist concentration, the serotonin concentration in each case was kept close to the EC_50_ value for wild-type (2 μM) and R64A (10 μM) (Fig 6c). Granisetron and palonosetron inhibition was measured at 1 nM; ondansetron and alosetron inhibition was measured at 0.1 nM (these concentrations where chosen to achieve a 50% inhibition for wild type upon co-application) (Fig 6d and 6e). Of note, co-application of setron in some cases has ∼100 fold lower effect than pre-application due to slow on-rates^41^. Mutational perturbation at Arg65 has a significant effect on inhibition by each setron, albeit to varying extents.

## Conclusion

A comprehensive structural analysis of multiple high-resolution structures of setron-bound 5-HT_3A_R complexes reveal several features of competitive antagonism that were not fully evident from the previous structural findings. Serotonin binds within a partially solvent-exposed cavity at the subunit interface and elicits Loop C closure, which may be coupled to channel opening. The setron-binding pocket, while involving overlapping residues, extends further into the complementary subunit. Setron-binding evokes varying degrees of Loop C closure and in some cases, almost to the same degree as a serotonin-bound state. The Loop C movements are associated with varying degrees of structural changes in the inner and outer β-strands that translate to small changes in the pore-lining M2 helices. Overall, setrons stabilize 5-HT_3A_R conformational states that are non-conductive, but appear to lie between the apo and serotonin-bound states. These findings therefore suggest that competitive antagonism in 5-HT_3A_R, and potentially in other pLGIC, may involve stabilizing intermediates along the activation pathway. With new emerging uses of setrons to treat psychiatric disorders, inflammation, substance abuse, and Alzheimer’s disease, these studies lay the foundation for the design of newer therapeutics with higher treatment efficacy and fewer off-target effects.

## Methods

### Electrophysiological measurements in oocytes

Mouse 5-HT_3A_R gene (purchased from GenScript) and mutant genes were inserted into pTLN plasmid. The plasmids were linearized with Mlu1 restriction enzyme by digesting overnight at 37 °C. The mMessage mMachine kit (Ambion) was used to make mRNA as per the manufacturer’s protocol and cleanup using RNAeasy kit (Qiagen). 3–10 ng of mRNA was injected into *X. laevis* oocytes (stages V–VI), and incubated for 2-5 days, after which current recordings were performed. Water injected oocytes were used as a control to verify that no endogenous currents were present. Female *X. laevis* were purchased from Nasco and kindly provided by W. F. Boron. Institutional Animal Care and Use Committee (IACUC) of Case Western Reserve University approved the animal experimental procedures. Oocytes were maintained in OR3 medium (GIBCO-BRL Leibovitz medium containing glutamate, 500 units each of penicillin and streptomycin, pH adjusted to 7.5, osmolarity adjusted to 197 mOsm) at 18 °C. Warner Instruments Oocyte Clamp OC-725 was used to perform two-electrode voltage-clamp experiments at a holding potential of −60 mV. Currents were sampled and digitized at 500 Hz with a Digidata 1332A. Clampfit 10.2 (Molecular Devices) was used to analyze experimental data. Perfusion solution consisted of 96mM NaCl, 2mM KCl, 1.8mM CaCl_2_, 1mM MgCl_2_, and 5mM HEPES (pH 7.4, osmolarity adjusted to 195 mOsM) was used at a flow rate of 6 ml/min. Chemical reagents (serotonin hydrochloride, alosetron hydrochloride, ondansetron hydrochloride, and palonosetron hydrochloride) were purchased from Sigma-Aldrich.

### Full-length 5-HT_3A_R cloning and transfection

The mouse 5-HT_3A_R (NCBI Reference Sequence: NM_001099644.1) gene was codon-optimized for *Spodoptera frugiperda* (Sf9) cells and purchased from GenScript. The construct consists of the 5-HT_3A_R gene along with a C-terminal 1D4-tag^42^ and four strep-tags (WSHPQFEK) at the N terminus, each separated by a linker sequence (GGGSGGGSGGGS) and followed by a TEV-cleavage sequence (ENLYFQG). Sf9 cells (Expression System) were grown in ESF921 medium (Expression Systems) at 28 °C without CO_2_ exchange and in absence of antibiotics. Cellfectin II reagent (Invitrogen) was used for transfection of recombinant 5-HT_3A_R bacmid DNA into sub-confluent Sf9 cells. After 72 h of transfection, the progeny 1 (P1) recombinant baculoviruses were obtained by collecting the cell culture supernatant. The P1 was then used to infect Sf9 cells which produced P2 viruses, and subsequently P3 viruses from the P2 virus stock. The P3 viruses were used for recombinant protein expression.

### 5-HT_3A_R expression and purification

Sf9 cells are grown to approximately 2.5 × 10^6^ per ml followed by infection with P3 viruses. After 72 h post-infection, the cells were centrifuged at 8,000g for 20 min at 4 °C to separate the supernatant from the pellet. The cell pellet was resuspended in 20 mM Tris-HCl, pH 7.5, 36.5 mM sucrose supplemented with 1% protease inhibitor cocktail (Sigma-Aldrich). Cells were sonicated on ice. Non-lysed cells were pelleted down by centrifugation (3,000g for 15 min) and the supernatant was collected. The membrane fraction was separated by ultracentrifugation (167,000g for 1 h) and solubilized in 50 mM Tris pH 7.5, 500 mM NaCl, 10% glycerol, 0.5% protease inhibitor and 1% C12E9 for 2 h at 4 °C. Non-solubilized material was removed by ultracentrifugation (167,000 g for 15 min). The solubilized membrane proteins containing 5-HT_3A_ receptors were bound with 1D4 beads pre-equilibrated with 20 mM HEPES pH 8.0, 150 mM NaCl and 0.01% C12E9 for 2 h at 4 °C. The non-bound proteins were removed by washing beads with 100 column volumes of 20 mM HEPES pH 8.0, 150 mM NaCl, and 0.01% C12E9 (buffer A). The protein was then eluted with 3 mg/ml 1D4 peptide (TETSQVAPA) which is solubilized in buffer A. Eluted protein was deglycosylated with PNGase F (NEB) by incubating 5 units of the enzyme per 1 μg of protein for 2 h at 37 °C under gentle agitation. Deglycosylated protein was then purified using a Superose 6 column (GE healthcare) equilibrated with buffer A. Purified protein was concentrated to 2–3 mg/ml using 50-kDa MWCO Millipore filters (Amicon) for cryo-EM studies.

### Cryo-EM sample preparation and data acquisition

5-HT_3A_R protein (∼2.5 mg/ml) was filtered and incubated with 100 μM drugs (Alosetron, Ondansetron, and Palonosetron) for 1 hour. Fluorinated Fos-choline-8 (Anatrace) was added to the protein sample to a final concentration of 3 mM. The protein was then blotted onto Cu 300 mesh Quantifoil 1.2/1.3 grids (Quantifoil Micro Tools) two times with 3.5 μl sample each time, and the grids were plunge frozen immediately into liquid ethane using a Vitrobot (FEI). The grids were imaged using a 300 kV FEI Titan Krios G3i microscope equipped with a Gatan K3 direct electron detector camera. Movies containing ∼50 frames were collected at 105,000× magnification (set on microscope) in super-resolution mode with a physical pixel size of 0.848 Å/pixel, dose per frame 1 e^-^/Å^2^. Defocus values of the images ranged from −1.0 to −2.5 µm (input range setting for data collection) as per the automated imaging software SerialEM^43^.

### Image processing

MotionCor^44^ was used to correct beam-induced motion using a B-factor of 150 pixels^2^. Super-resolution images were binned (2×2) in Fourier space, making a final pixel size of 0.848 Å. Entire data processing was conducted in RELION 3.1^45^. CTF of the motion-corrected micrographs were estimated using Gctf software^46^. Auto-picked particles from total micrographs (Table 1) from individual datasets (each drug) were subjected to 2D classification to remove suboptimal particles. An initial 3D reference model was generated from the 5-HT_3A_R-apo cryo-EM structure (RCSB Protein Data Bank code (PDB ID): 6BE1). The model was low-pass filtered at 60 Å using EMAN2^47^. Iterative 3D classifications, 3D auto-refinements, and bayesian polishing generated density model of Alosetron, Ondansetron and Palonosetron bound 5-HT_3A_R with 42, 065 particles, 67, 333 particles, and 91,163 particles, respectively. Per-particle contrast transfer function (CTF) refinement and beam tilt correction were applied followed by a final 3D-autorefinement. A soft mask was generated in RELION and used during the post-processing step, which resulted in an overall resolution of 2.92 Å, 3.06 Å, and 3.32 Å for 5-HT_3A_R-Alo, 5-HT_3A_R-Ondan and 5-HT_3A_R-Palono, respectively (calculated based on the gold-standard Fourier shell coefficient (FSC) = 0.143 criterion, Table 1). B-factor estimation and map sharpening were performed in the post-processing step in RELION. The ResMap program was used to calculate local resolutions ^48^.

**Table 1:** Cryo-EM data collection/processing.

|  | 5-HT<sub>3A</sub>-Alosetron<br>(EMDB-21511; PDB-6W1J) | 5-HT<sub>3A</sub>-Ondansetron<br>(EMDB-21512; PDB-6W1M) | 5-HT<sub>3A</sub>-Palonosetron<br>(EMDB-21518; PDB-6W1Y) |
| --- | --- | --- | --- |
| **Data collection and processing** |  |  |  |
| Magnification |  | 105,000x |  |
| Voltage (kV) |  | 300 |  |
| Data collection mode |  | Super-resolution |  |
| Electron exposure (e-/Å<sup>2</sup>) |  | 50 |  |
| Defocus range (μm) |  | -1.2 to -2.5 |  |
| Physical Pixel size (Å/pixel) |  | 0.848 |  |
| Symmetry imposed |  | C5 |  |
| Initial particle images (no.) | 568,452 | 449,628 | 1,114,542 |
| Final particle images (no.) | 42,065 | 67,333 | 91,163 |
| Map resolution (unmasked, Å) at<br>FSC<sub>143</sub> | 3.2 | 3.4 | 3.7 |
| Map resolution (masked, Å) at<br>FSC<sub>143</sub> | 2.92 | 3.06 | 3.32 |
| Map resolution range (Local<br>resolution) | 2.5-4.5 | 2.5-4.5 | 2.5-4.5 |
| **Refinement** |  |  |  |
| Initial model used (PDB code) | 6BE1 | 6BE1 | 6BE1 |
| Map sharpening *B* factor (Å<sup>2</sup>) | -30 | -30 | -70 |
| Model composition |  |  |  |
| Non-hydrogen atoms | 16,861 | 16,885 | 16,885 |
| Protein residue numbers | 393 | 394 | 394 |
| Ligand atoms | 586 | 585 | 585 |
| *B* factors (Å<sup>2</sup>) |  |  |  |
| Protein | 101.61 | 115.04 | 132.89 |
| Ligand | 103.61 | 100.96 | 116.04 |
| R.m.s. deviations |  |  |  |
| Bond lengths (Å) | 0.008 | 0.008 | 0.009 |
| Bond angles (°) | 0.910 | 0.991 | 1.069 |
| Validation |  |  |  |
| MolProbity score | 1.38 (97<sup>th</sup> Percentile) | 1.48 (96<sup>th</sup> Percentile) | 1.41 (97<sup>th</sup> Percentile) |
| Clashscore | 2.22 (99<sup>th</sup> Percentile) | 3.19 (97<sup>th</sup> Percentile) | 2.53 (98<sup>th</sup> Percentile) |
| Poor rotamers (%) | 0.82 | 0.27 | 0.82 |
| Ramachandran plot |  |  |  |
| Favored (%) | 94.40 | 94.62 | 94.62 |
| Allowed (%) | 5.60 | 5.38 | 5.38 |
| Disallowed (%) | 0.00 | 0.00 | 0.00 |

### 5-HT_3A_R model building

The final refined models have clear density of residues Thr7–Leu335 and Leu397–Ser462. The unobserved density at the region of (336–396) is comprised of an unstructured loop which links the amphipathic MX helix and the MA helix. The 5-HT_3A_R-apo cryo-EM structure (PDB ID: 6BE1) was used as an initial model and refined against its EM-derived map using PHENIX software package ^49^, using rigid body, local grid, NCS, and gradient minimization parameters. COOT is used for manual model building ^50^. Real space refinement in PHENIX yielded the final model with a final model to map cross-correlation coefficient of 0.834 (5-HT_3A_R-Palono), 0.846 (HT_3A_R-Ondan), and 0.848 (5-HT_3A_R-Alo). Stereochemical properties of the model were validated by Molprobity^51^. The pore profile was calculated using the HOLE program^52^. Figures were prepared using PyMOL v.2.0.4 (Schrödinger, LLC).

### MD Simulation Setup and Protocol

The cryo-EM-derived structures of palonosetron-bound, alosetron-bound, and ondansetron-bound 5-HT_3A_R pentamer were prepared for MD simulations with the Protein Prep Wizard in the SchrÖdinger scientific software suite 2019-2 using default settings (Small-Molecule Drug Discovery Suite 2019-2, Schrödinger, LLC, New York, NY, 2019). This protocol adds missing hydrogen atoms to the initial protein-ligand complex. After the initial preparatory steps and protonation assignment of side chains, a brief restrained energy minimization *in vacuo* using the OPLS3 force field^53^ was carried out to finalize system setup for each protein-ligand complex. Each setron-5-HT_3A_R complex was then embedded into a 1-palmitoyl-2-oleoyl phosphatidyl choline (POPC) bilayer using the Membrane Builder tool of the CHARMM-GUI webserver ^54^. The system was then solvated with TIP3P water and 150 mM NaCl were added to the simulation system by replacing random water molecules. Excess sodium ions were added to neutralize the charge of each protein-ligand complex. The resulting simulation systems had initial dimensions of ∼130×130×207 Å^3^ and consisted of the 5-HT_3A_R pentamer, the setrons bound to each 5-HT_3A_R subunit, ∼400 POPC molecules, ∼83,000 water molecules, ∼240 sodium ions, and ∼220 chloride ions, for a total of ∼330,000-346,000 atoms. Throughout this work we reference data from our previously published simulation of granisetron-bound 5-HT_3A_R^26^ in comparison to simulations of these three novel setron-bound 5-HT_3A_R complexes.

The CHARMM36m forcefield^55^ was used to parameterize the protein and lipid atoms within each simulation system. Initial parameters for palonosetron, alosetron, and ondansetron were obtained from the ParamChem webserver using the CHARMM general force field ^56^ (https://cgenff.parmchem.org). Parameters were validated according to the procedure described^56^. Said validation required quantum calculations performed with Gaussian 16 (Gaussian 16, Revision C.01, Gaussian, Inc., Wallingford CT, 2016) to finalize the charges and dihedrals defined within our setron molecule models.

MD simulations were run using GROMACS 2018.6^57^software with a timestep of 2 fs, following a steepest descent energy minimization run for 5000 steps, as well as 100 ps isothermal-isovolumetric (NVT) and 52 ns isothermal-isobaric (NPT) equilibration runs. The NVT equilibration was performed to initially heat the model systems after the steepest descents minimization. This step was performed with restraints on protein, membrane, and ligand molecule heavy atoms relative to their starting conformation. The NPT equilibration runs were performed in 5 steps of 10 ns each, within which the system was allowed to relax with gradually released restraints until finally the system was allowed to equilibrate for 2 ns of unrestrained NPT equilibration. This was followed by a 100 ns production run in isothermal-isobaric conditions. System temperature and pressure were maintained at 300 K and 1 bar, respectively, using velocity rescale^58^ for temperature coupling and Parrinello-Rahman barostat for pressure coupling during equilibration. Semi-isotropic pressure coupling and the Nose-Hoover thermostat^59^ were applied during production runs. All bonds involving hydrogens were constrained using the LINCS algorithm^60^. Short-range nonbonded interactions were cut at 12 Å. Long-range electrostatic interactions were computed using the Particle Mesh Ewald summation with a Fourier grid spacing of 1.2 Å.

Trajectory analyses were performed using a combination of Visual Molecular Dynamics (VMD)^61^ and the GROMACS analysis toolkit^62^ over equidistant frames of our production simulations using a 500 ps stride. In particular, all RMSD measurements and loop C orientations were obtained after aligning simulation frames onto the coordinates of the initial cryo-EM structure by comparing Cα atoms in the helices and β-sheets of the ECD. RMSD calculations were assessed for each setron by evaluating the difference in heavy atoms of the ligands between each simulation frame and the initial cryo-EM structure conformation. Similarly, Loop C RMSD’s were calculated by comparing the Cα, backbone carbonyl carbon, and backbone nitrogen atoms of residues 200 through 205 relative to their conformation in the initial cryo-EM resolved structures. To measure the orientation of Loop C, we defined a custom Loop C dihedral as being drawn between the alpha carbons of residues Ala208, Phe199, Glu198, and Ile203. To determine whether Loop C adopted a ‘closed’ or ‘open’ conformation we evaluated the distance between the Arg65 and Asp202 side chains, measured by a minimum distance of their respective polar side chain atoms for each analysed simulation frame. To evaluate how well solvated the setron binding sites were throughout our simulations, we counted the number of water oxygen atoms within 3 Å of any setron atoms for each simulation frame. Structural interaction fingerprints were calculated with an in-house python script that monitored 5-HT_3A_R interactions with each setron. Specifically, for each residue of 5-HT_3A_R, setron-protein interactions were calculated as a 9-bit representation based on the following 9 types of interactions: apolar (van der Waals), face-to-face aromatic, edge-to-face aromatic, hydrogen-bond interactions with the protein either as a donor or acceptor, electrostatic with either the protein acting as a positive or negative charge, one-water-mediated hydrogen bond, and two-water-mediated hydrogen bonds. A distance cutoff of 4.5 Å was used to identify apolar interactions between two non-polar atoms (carbon atoms), while a cutoff of 4 Å was used to evaluate aromatic and electrostatic interactions. Residue interactions were evaluated for protein side chain atoms only. Average probabilities and errors for each interaction type were estimated using a two-state Markov model, sampling the transition matrix posterior distribution using standard Dirichlet priors for the transition probabilities^63^.

### Data Availability Accession Numbers

The coordinates of the 5-HT_3A_R-drugs structure and the Cryo-EM map have been deposited in wwPDB and EMDB with following accession number. PDB ID: 6W1J; EMBD ID: EMD-21511 for 5-HT_3A_R-Alo, PDB ID: 6W1M; EMBD ID: EMD-21512 for 5-HT_3A_R-Ondan and PDB ID: 6W1Y; EMBD ID: EMD-21518 for 5-HT_3A_R-Palono. All relevant data are available from the corresponding author upon reasonable request.

## Acknowledgements

We acknowledge the use of instruments at the Cryo-Electron Microscopy Core at the CWRU School of Medicine. We are grateful to Dr. Kunpeng Li for assistance with cryo-EM imaging and data collection. We thank Denice Major for assistance with hybridoma and cell culture at the Department of Ophthalmology and Visual Sciences (supported by the National Institutes of Health Core Grant P30EY11373). We thank Dr. Walter F. Boron for kindly providing us *Xenopus* oocytes and for unrestricted access of the oocyte rig. We are deeply appreciative of the support provided by Dr. Fraser Moss and Mr. Brian Zeise with the oocyte rig. We are very grateful to the members of the Chakrapani lab for critical reading and comments on the manuscript. Computations were run on resources available through the Scientific Computing Facility at the Icahn School of Medicine at Mount Sinai and the Extreme Science and Engineering Discovery Environment under MCB080077 (to M.F.), which is supported by National Science Foundation grant number ACI-1548562. This work was supported by the National Institutes of Health grants R01GM108921, R01GM131216, R35GM134896, and Cryo-EM supplements: 3R01GM108921-03S1, R01GM108921-5S1, 3R01GM131216-1S1 to S.C and the AHA postdoctoral Fellowship to A.K (20POST35210394) and S.B (17POST33671152).

## Author Contributions

S.B and S.C conceived the project and designed experimental procedures. S.B purified the protein, optimized the cryo-EM sample preparation, carried out data collection, analysis, model building, refinement, and performed two-electrode voltage-clamp recordings. A. Kumar contributed to cryo-EM data collection and functional analysis. S.C supervised the execution of the experiments, data analysis, and interpretation. E.G contributed to analysis and mapping of structural differences using MATLAB. S.R, with the assistance of A. Kapoor, performed the MD simulations, docking, and other computational analyses under the supervision of M.F. All authors contributed to manuscript preparation.

## Competing Financial Interest

The authors declare that there is no competing financial interest.

## Figure Legends

**Supplemental Figure 1.**
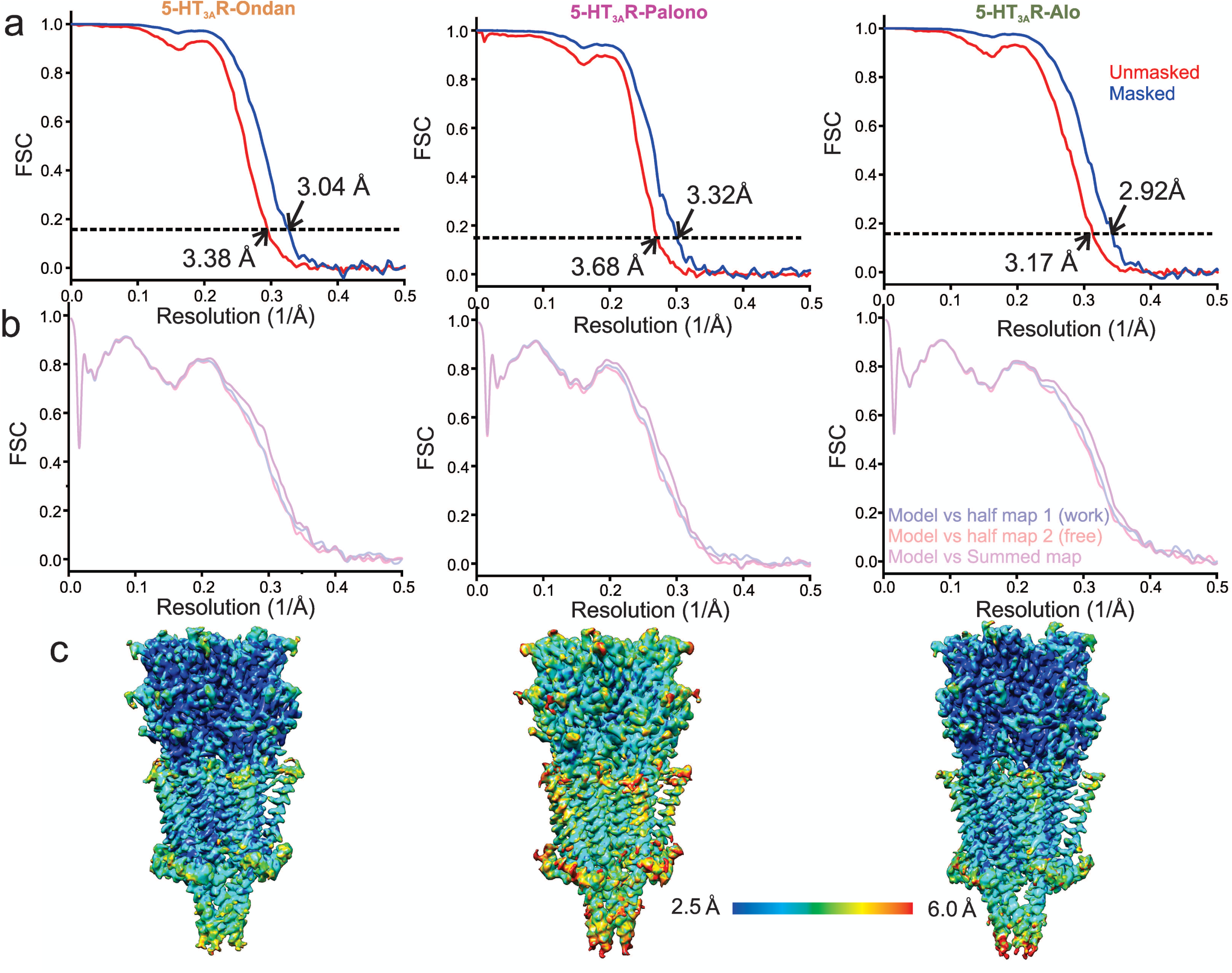
Resolution estimation and validation of Cryo-EM models. **a** For 5-HT_3A_R-Ondan (*left*), 5-HT_3A_R-Palono (*middle*), and 5-HT_3A_R-Alo (*right*) structures, gold standard Fourier Shell Correlation (FSC) curves generated by comparing two independent half maps produced during refinement in RELION. Horizontal dotted line represents the FSC threshold at 0.143. Nominal resolution of setron-bound 5-HT_3A_R at FSC_0.143_ of unmasked (red FSC curve) and masked (blue FSC curve) maps are indicated by arrow. **b** Model and Cryo-EM map correlation indicates no model bias in 5-HT_3A_R. **c** Side-view of setron-5-HT_3A_R maps colored based on the local resolution.

**Supplemental Figure 2.**
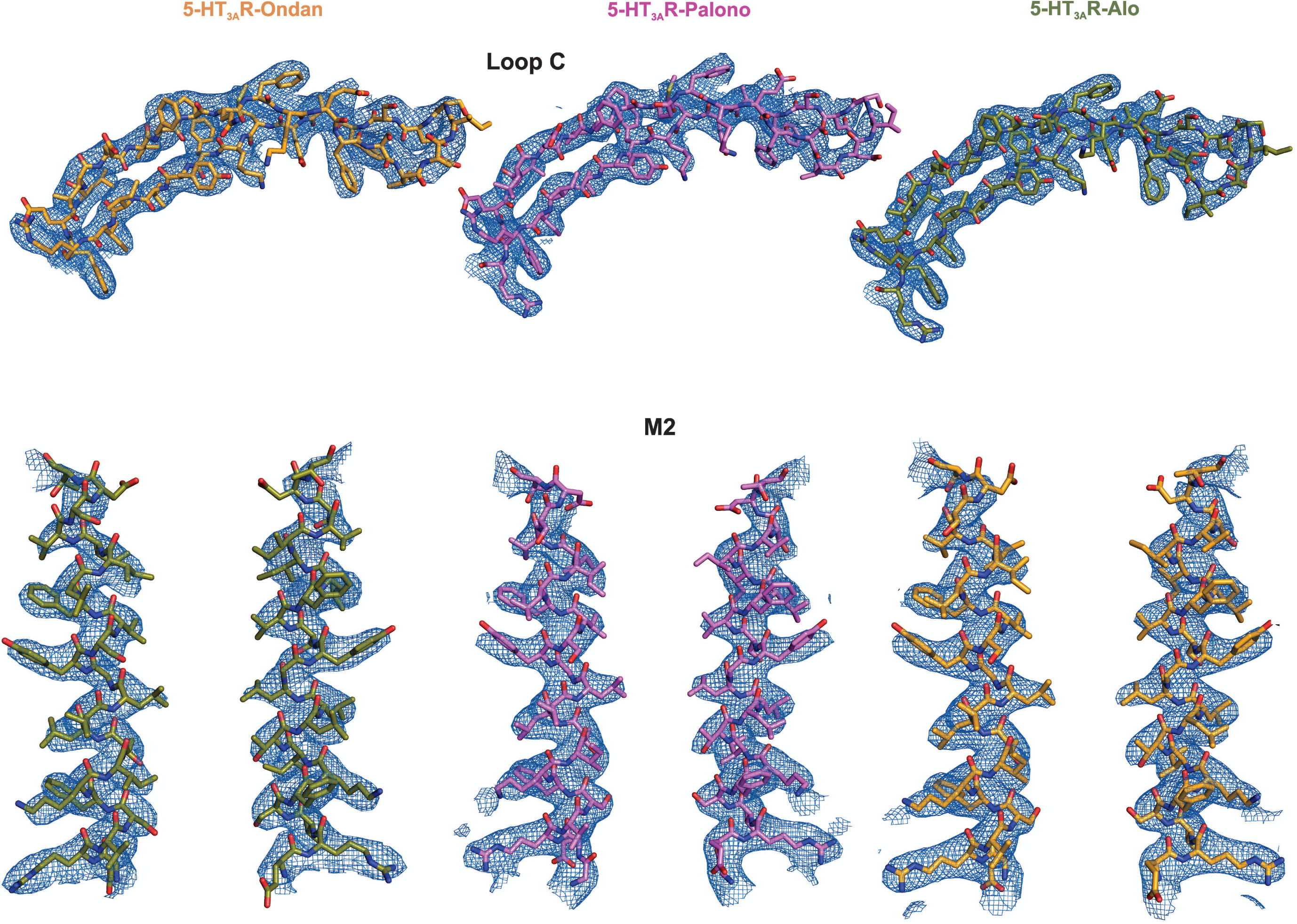
Assessment of Cryo-EM map quality and model fitting in the map. Validation of Loop C modelling in the setron-5-HT_3A_R maps (*top*). Residues are represented in stick and densities are shown in blue mesh. Loop C meshes are contoured as follows: 5-HT_3A_R-Ondan (*left*) at 6 σ, 5-HT_3A_R-Palono (*middle*) at 7 σ, and 5-HT_3A_R-Alo (*right*) at 6 σ. Cryo-EM densities are shown for the M2 helices (*bottom*; two subunits are shown for clarity). Contour level of the mesh for 5-HT_3A_R-Ondan (*left*), 5-HT_3A_R-Palono (*middle*), and 5-HT_3A_R-Alo are each at 5σ.

**Supplemental Figure 3.**
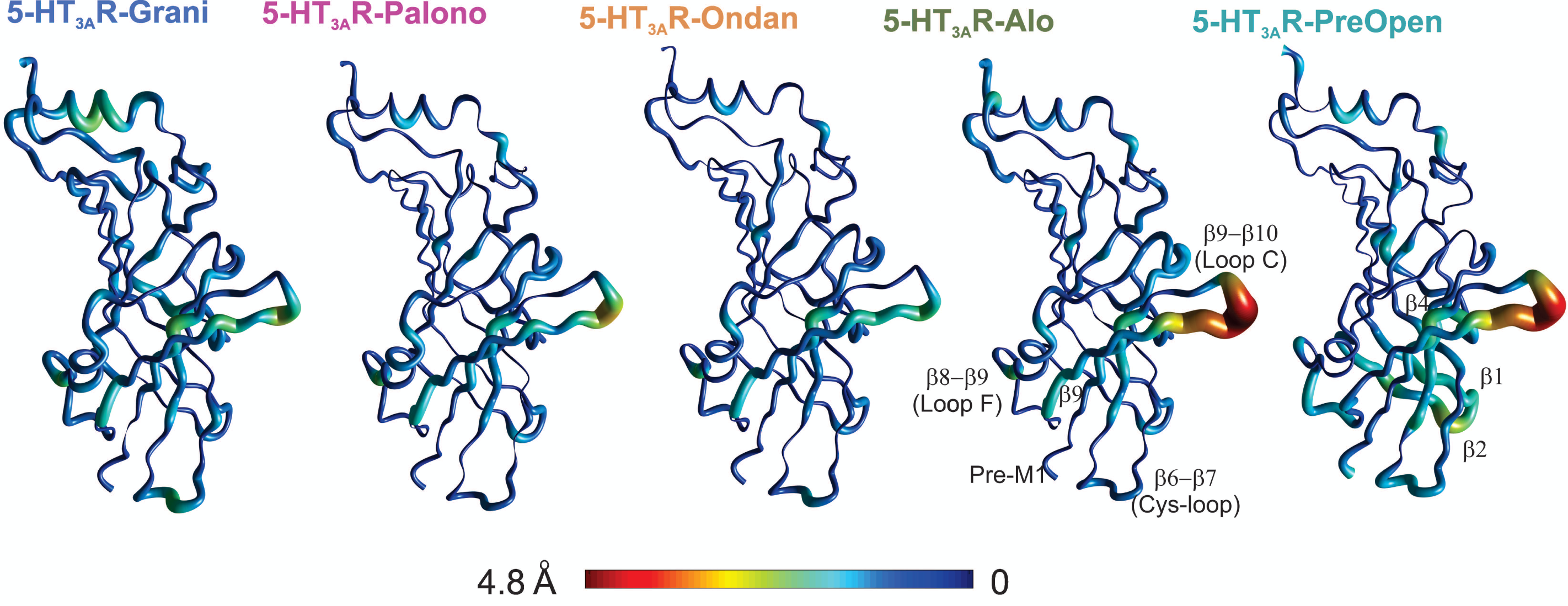
Conformational differences in the ECD of setron- and serotonin-bound structures compared to Apo-5-HT_3A_R. Pentameric assembly of setron- and serotonin-bound structures were aligned to Apo-5-HT_3A_R. A cubic spline interpolation was then done to smoothly connect cα displacement for each structure and mapped by short cylinders, whose diameters are equivalent to the displacement at that position compared to Apo-5-HT_3A_R. The color was also scaled to the same value using the color map shown. The analysis was done in Matlab v2019a (Mathworks, Natick MA).

**Supplemental Figure 4.**
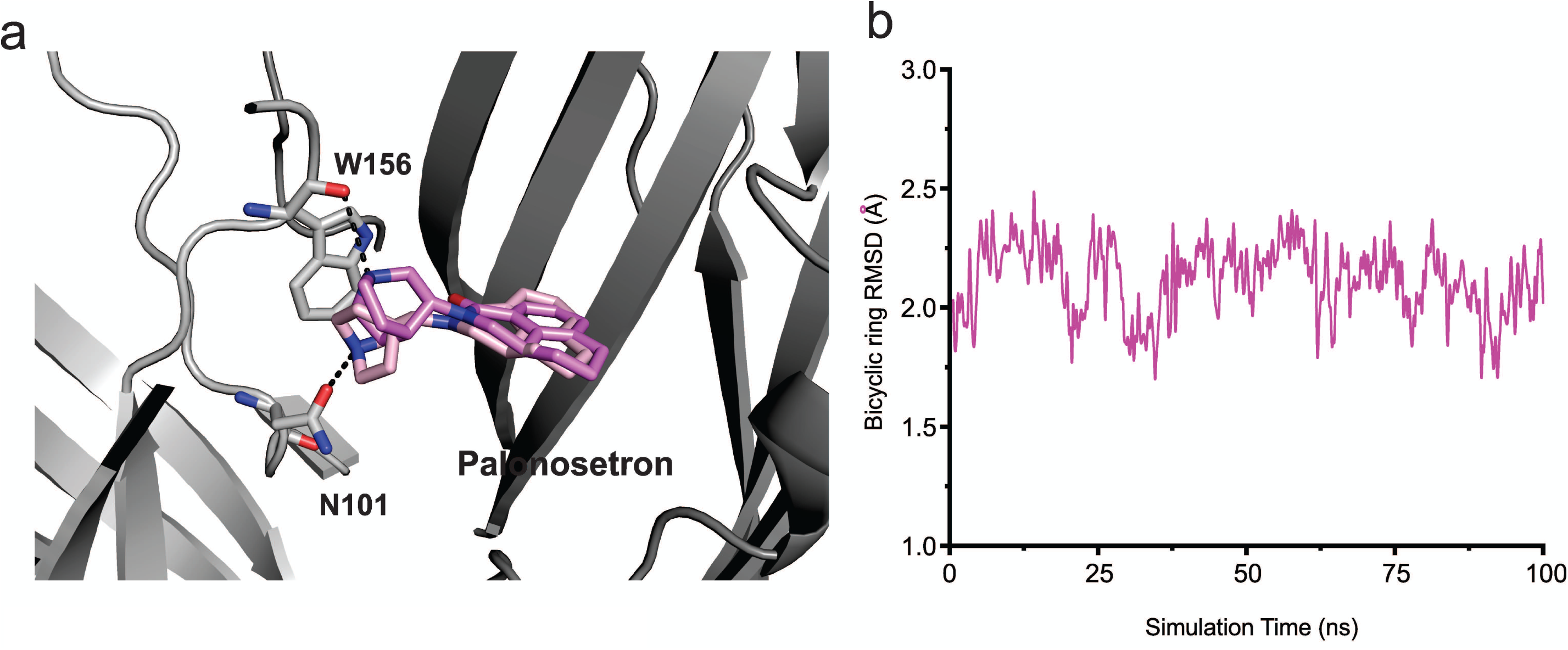
Palonosetron orientation in the MD simulation. **a** Representative views of various palonosetron orientations during the 100 ns simulation. When the tertiary amine nitrogen in the bicyclic ring is pointing up, it interacts with the carbonyl oxygen of Trp156 and when it points down, it interacts with carbonyl oxygen of Asn101 side chain. **b** Time evolution of Root Mean Square Deviation (RMSD) of the bicyclic ring to its initial cryo-EM position.

**Supplemental Figure 5.**
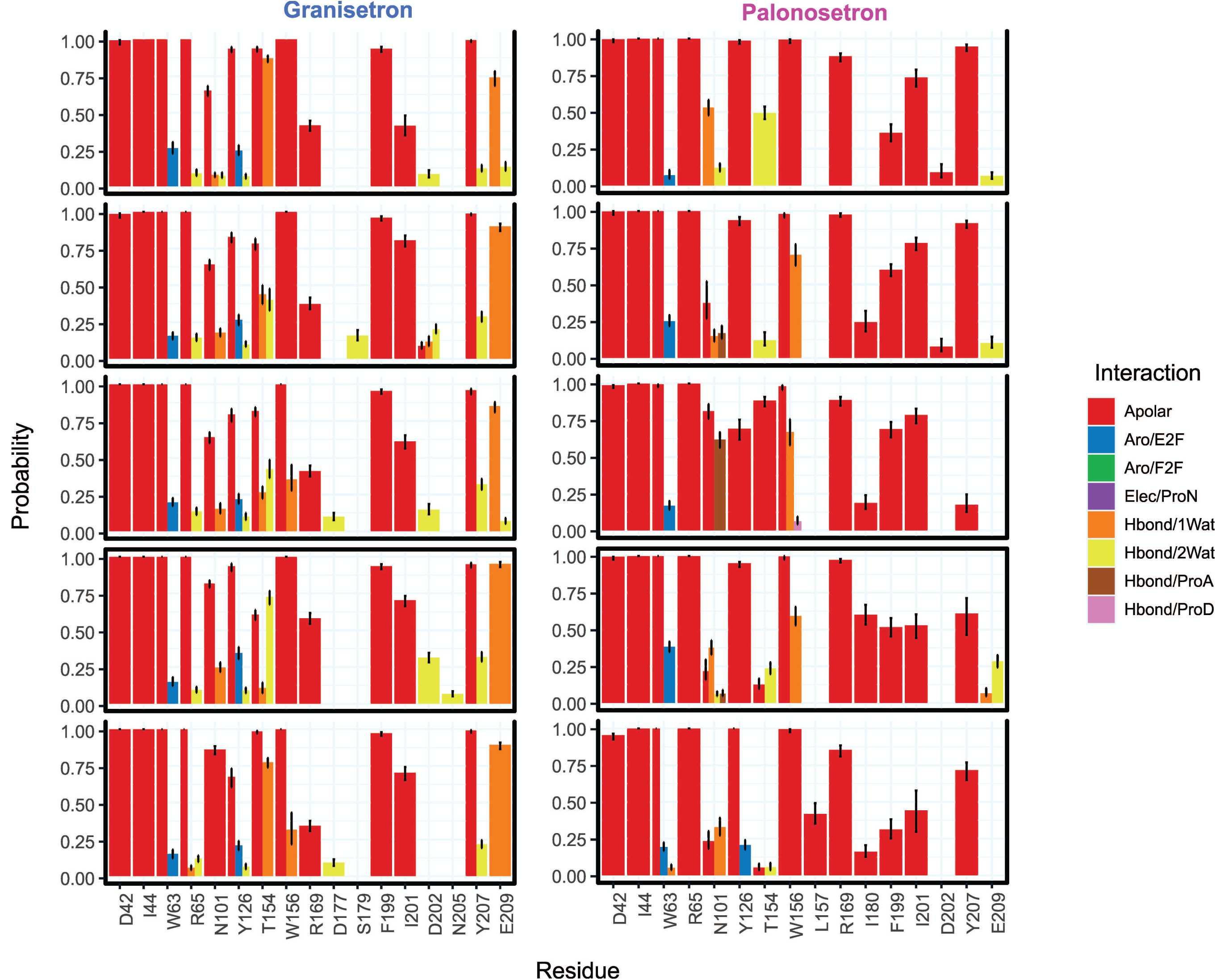
Interaction Fingerprints in 5-HT_3A_R-Grani and 5-HT_3A_R-Palono structures. Molecular interactions between setron and 5-HT_3A_R in 5-HT_3A_R-Grani and 5-HT_3A_R-Palono structures during MD simulation. 5-HT_3A_R-setron interaction fingerprints were derived for each ligand-protein complex and each protomer chain from 100 ns MD simulations. Interactions were assessed between ligand and protein side chain atoms and reported for those that had an average probability above 5%. Nine interaction types were calculated: apolar (hydrophobic), face- to-face aromatic (Aro_F2F), edge-to-face aromatic (Aro_E2F), hydrogen bond with the protein as hydrogen bond donor (Hbond_ProD), hydrogen bond with the protein as hydrogen bond acceptor (Hbond_ProA), electrostatic with the protein positively charged (Elec_ProP), electrostatic with the protein negatively charged (Elec_ProN), one-water-mediated and two-water-mediated hydrogen bond interactions (Hbond_1Wat and Hbond_2Wat). Electrostatic interactions with positively charged protein residues (Elec_ProP) were not found in any of our studied setron-bound 5-HT_3A_R simulations.

**Supplemental Figure 6.**
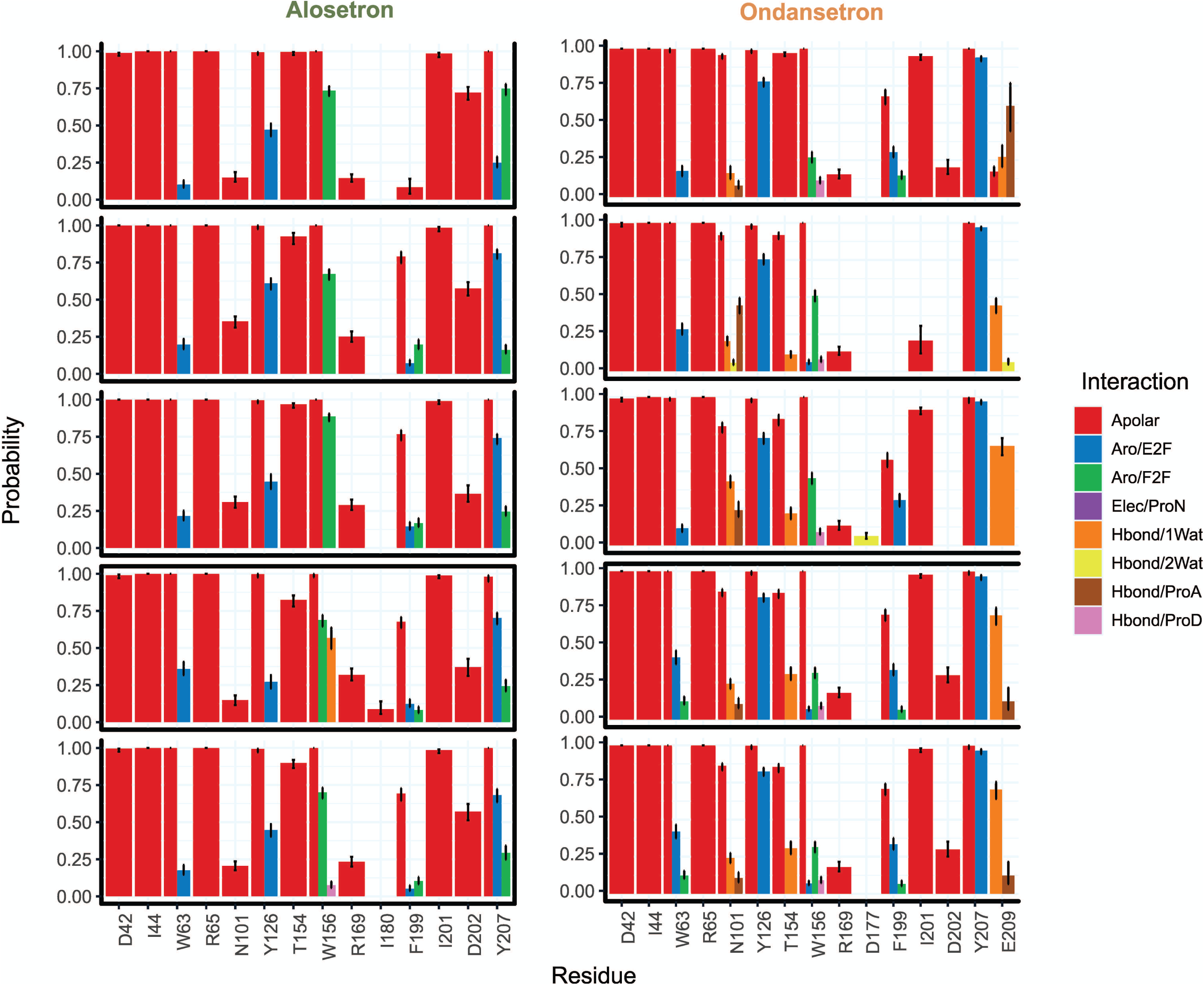
Interaction Fingerprints in 5-HT_3A_R-Ondan and 5-HT_3A_R-Alo structures. Molecular interactions between setron and 5-HT_3A_R in 5-HT_3A_R-Ondan and 5-HT_3A_R-Alo structures during MD simulation. The interaction color codes are as described in the figure legends for Supplemental Figure 5.

**Supplemental Figure 7.**
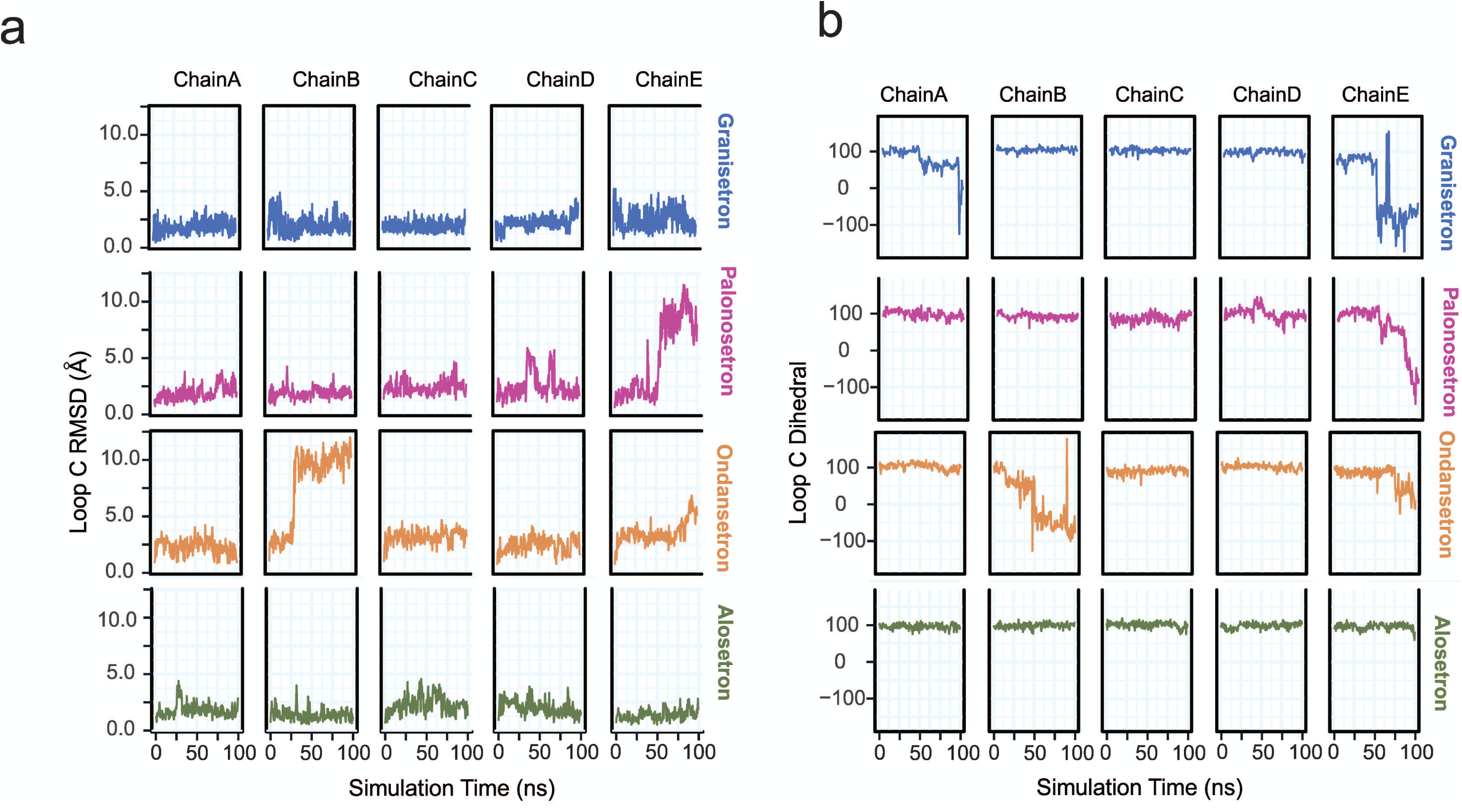
Conformational differences in Loop C. **a** Time evolution of loop C RMSD evaluated by quantifying the distance of Cα, carbonyl carbon, and backbone nitrogen atoms of residues Ser200 to Asn205 relative to their initial cryo-EM conformations separated by protomer chain. **b** Time evolution of loop C custom dihedral (defined as a dihedral drawn from Cα atoms of residues Ala208, Phe199, Glu198, and Ile203). Loop C is oriented away from the binding site when this dihedral is measured to be large and is oriented toward the binding site when this dihedral is low or negative.

## References

1 Schworer, H., Racke, K. & Kilbinger, H. Cisplatin increases the release of 5-hydroxytryptamine (5-HT) from the isolated vascularly perfused small intestine of the guinea-pig: involvement of 5-HT3 receptors. Naunyn Schmiedebergs Arch Pharmacol 344, 143–149, doi:10.1007/bf00167211 (1991).

2 Gilmore, J., D’Amato, S., Griffith, N. & Schwartzberg, L. Recent advances in antiemetics: new formulations of 5HT3-receptor antagonists. Cancer Manag Res 10, 1827–1857, doi:10.2147/CMAR.S166912 (2018).

3 Spiller, R. C. Targeting the 5-HT3 receptor in the treatment of irritable bowel syndrome. Current Opinion in Pharmacology 11, 68–74, doi: https://doi.org/10.1016/j.coph.2011.02.005 (2011).

4 Hsu, E. S. A review of granisetron, 5-hydroxytryptamine3 receptor antagonists, and other antiemetics. Am J Ther 17, 476–486, doi:10.1097/MJT.0b013e3181ea7821 (2010).

5 de Wit, R., Aapro, M. & Blower, P. R. Is there a pharmacological basis for differences in 5-HT3-receptor antagonist efficacy in refractory patients? Cancer Chemother Pharmacol 56, 231–238, doi:10.1007/s00280-005-1033-0 (2005).

6 Friedel, D., Thomas, R. & Fisher, R. S. Ischemic colitis during treatment with alosetron. Gastroenterology 120, 557–560, doi:10.1053/gast.2001.21177 (2001).

7 Engel, M., Smidt, M. & Van Hooft, J. The serotonin 5-HT3 receptor: a novel neurodevelopmental target. Frontiers in Cellular Neuroscience 7, doi:10.3389/fncel.2013.00076 (2013).

8 Lummis, S. C. R. 5-HT3 Receptors. Journal of Biological Chemistry 287, 40239–40245, doi:10.1074/jbc.R112.406496 (2012).

9 Kia, H. K. et al. Localization of 5-HT3 receptors in the rat spinal cord: immunohistochemistry and in situ hybridization. Neuroreport 6, 257–261 (1995).

10 Bétry, C. et al. Role of 5-HT(3) Receptors in the Antidepressant Response. Pharmaceuticals 4, 603–629, doi:10.3390/ph4040603 (2011).

11 Thompson, A. J. & Lummis, S. C. 5-HT3 receptors. Curr Pharm Des 12, 3615–3630 (2006).

12 Gershon, M. D. Review article: serotonin receptors and transporters -- roles in normal and abnormal gastrointestinal motility. Alimentary pharmacology & therapeutics 20 Suppl 7, 3–14, doi:10.1111/j.1365-2036.2004.02180.x (2004).

13 Maricq, A. V., Peterson, A. S., Brake, A. J., Myers, R. M. & Julius, D. Primary structure and functional expression of the 5HT3 receptor, a serotonin-gated ion channel. Science 254, 432, doi:10.1126/science.1718042 (1991).

14 Niesler, B. et al. Characterization of the novel human serotonin receptor subunits 5-HT3C,5-HT3D, and 5-HT3E. Mol Pharmacol 72, 8–17, doi:10.1124/mol.106.032144 (2007).

15 Davies, P. A. et al. The 5-HT3B subunit is a major determinant of serotonin-receptor function. Nature 397, 359–363, doi:10.1038/16941 (1999).

16 Kelley, S. P., Dunlop, J. I., Kirkness, E. F., Lambert, J. J. & Peters, J. A. A cytoplasmic region determines single-channel conductance in 5-HT3 receptors. Nature 424, 321–324, doi:10.1038/nature01788 (2003).

17 Thompson, A. J. & Lummis, S. C. Discriminating between 5-HT(3)A and 5-HT(3)AB receptors. Br J Pharmacol 169, 736–747, doi:10.1111/bph.12166 (2013).

18 Hammer, C. et al. Replication of functional serotonin receptor type 3A and B variants in bipolar affective disorder: a European multicenter study. Transl Psychiatry 2, e103, doi:10.1038/tp.2012.30 (2012).

19 Gregory, R. E. & Ettinger, D. S. 5-HT3 receptor antagonists for the prevention of chemotherapy-induced nausea and vomiting. A comparison of their pharmacology and clinical efficacy. Drugs 55, 173–189, doi:10.2165/00003495-199855020-00002 (1998).

20 Kovac, A. L. Comparative Pharmacology and Guide to the Use of the Serotonin 5-HT3 Receptor Antagonists for Postoperative Nausea and Vomiting. Drugs 76, 1719–1735, doi:10.1007/s40265-016-0663-3 (2016).

21 Smith, H. S., Cox, L. R. & Smith, E. J. 5-HT3 receptor antagonists for the treatment of nausea/vomiting. Ann Palliat Med 1, 115–120, doi:10.3978/j.issn.2224-5820.2012.07.07 (2012).

22 Kesters, D. et al. Structural basis of ligand recognition in 5-HT3 receptors. EMBO Rep 14, 49–56, doi:10.1038/embor.2012.189 (2013).

23 Hibbs, R. E. et al. Structural determinants for interaction of partial agonists with acetylcholine binding protein and neuronal alpha7 nicotinic acetylcholine receptor. Embo J 28, 3040–3051 (2009).

24 Price, K. L., Lillestol, R. K., Ulens, C. & Lummis, S. C. Palonosetron-5-HT3 Receptor Interactions As Shown by a Binding Protein Cocrystal Structure. ACS Chem Neurosci 7, 1641–1646, doi:10.1021/acschemneuro.6b00132 (2016).

25 Polovinkin, L. et al. Conformational transitions of the serotonin 5-HT3 receptor. Nature 563, 275–279, doi:10.1038/s41586-018-0672-3 (2018).

26 Basak, S. et al. Molecular mechanism of setron-mediated inhibition of full-length 5-HT3A receptor. Nat Commun 10, 3225, doi:10.1038/s41467-019-11142-8 (2019).

27 Hassaine, G. et al. X-ray structure of the mouse serotonin 5-HT3 receptor. Nature 512, 276–281, doi:10.1038/nature13552 (2014).

28 Basak, S. et al. Cryo-EM structure of 5-HT3A receptor in its resting conformation. Nat Commun 9, 514, doi:10.1038/s41467-018-02997-4 (2018).

29 Basak, S., Gicheru, Y., Rao, S., Sansom, M. S. P. & Chakrapani, S. Cryo-EM reveals two distinct serotonin-bound conformations of full-length 5-HT3A receptor. Nature 563, 270–274, doi:10.1038/s41586-018-0660-7 (2018).

30 Del Cadia, M. et al. Exploring a potential palonosetron allosteric binding site in the 5-HT(3) receptor. Bioorg Med Chem 21, 7523–7528, doi:10.1016/j.bmc.2013.09.028 (2013).

31 Yan, D., Schulte, M. K., Bloom, K. E. & White, M. M. Structural features of the ligand-binding domain of the serotonin 5HT3 receptor. The Journal of biological chemistry 274, 5537–5541 (1999).

32 Duffy, N. H., Lester, H. A. & Dougherty, D. A. Ondansetron and granisetron binding orientation in the 5-HT(3) receptor determined by unnatural amino acid mutagenesis. ACS Chem Biol 7, 1738–1745, doi:10.1021/cb300246j (2012).

33 Thompson, A. J. et al. Locating an antagonist in the 5-HT3 receptor binding site using modeling and radioligand binding. J Biol Chem 280, 20476–20482, doi:10.1074/jbc.M413610200 (2005).

34 Beene, D. L. et al. Cation-pi interactions in ligand recognition by serotonergic (5-HT3A) and nicotinic acetylcholine receptors: the anomalous binding properties of nicotine. Biochemistry 41, 10262–10269 (2002).

35 Thompson, A. J., Padgett, C. L. & Lummis, S. C. Mutagenesis and molecular modeling reveal the importance of the 5-HT3 receptor F-loop. J Biol Chem 281, 16576–16582 (2006).

36 Hansen, S. B. et al. Structures of Aplysia AChBP complexes with nicotinic agonists and antagonists reveal distinctive binding interfaces and conformations. Embo J 24, 3635–3646 (2005).

37 Purohit, P. & Auerbach, A. Loop C and the mechanism of acetylcholine receptor-channel gating. The Journal of general physiology 141, 467–478, doi:10.1085/jgp.201210946 (2013).

38 Marcus, Y. Ionic radii in aqueous solutions. Chemical Reviews 88, 1475–1498, doi:10.1021/cr00090a003 (1988).

39 Guros, N. B., Balijepalli, A. & Klauda, J. B. Microsecond-timescale simulations suggest 5-HT-mediated preactivation of the 5-HT3A serotonin receptor. Proceedings of the National Academy of Sciences of the United States of America 117, 405–414, doi:10.1073/pnas.1908848117 (2020).

40 Ruepp, M. D., Wei, H., Leuenberger, M., Lochner, M. & Thompson, A. J. The binding orientations of structurally-related ligands can differ; A cautionary note. Neuropharmacology 119, 48–61, doi:10.1016/j.neuropharm.2017.01.023 (2017).

41 Lummis, S. C. & Thompson, A. J. Agonists and antagonists induce different palonosetron dissociation rates in 5-HT(3)A and 5-HT(3)AB receptors. Neuropharmacology 73, 241–246, doi:10.1016/j.neuropharm.2013.05.010 (2013).

42 MacKenzie, D., Arendt, A., Hargrave, P., McDowell, J. H. & Molday, R. S. Localization of binding sites for carboxyl terminal specific anti-rhodopsin monoclonal antibodies using synthetic peptides. Biochemistry 23, 6544–6549 (1984).

43 Mastronarde, D. N. Automated electron microscope tomography using robust prediction of specimen movements. J Struct Biol 152, 36–51, doi:10.1016/j.jsb.2005.07.007 (2005).

44 Zheng, S. Q. et al. MotionCor2: anisotropic correction of beam-induced motion for improved cryo-electron microscopy. Nat Methods 14, 331–332, doi:10.1038/nmeth.4193 (2017).

45 Fernandez-Leiro, R. & Scheres, S. H. W. A pipeline approach to single-particle processing in RELION. Acta Crystallogr D Struct Biol 73, 496–502, doi:10.1107/S2059798316019276 (2017).

46 Mindell, J. A. & Grigorieff, N. Accurate determination of local defocus and specimen tilt in electron microscopy. J Struct Biol 142, 334–347 (2003).

47 Tang, G. et al. EMAN2: an extensible image processing suite for electron microscopy. J Struct Biol 157, 38–46, doi:10.1016/j.jsb.2006.05.009 (2007).

48 Kucukelbir, A., Sigworth, F. J. & Tagare, H. D. Quantifying the local resolution of cryo-EM density maps. Nat Methods 11, 63–65, doi:10.1038/nmeth.2727 (2014).

49 Adams, P. D. et al. PHENIX: building new software for automated crystallographic structure determination. Acta crystallographica. Section D, Biological crystallography 58, 1948–1954 (2002).

50 Emsley, P. & Cowtan, K. Coot: model-building tools for molecular graphics. Acta crystallographica. Section D, Biological crystallography 60, 2126–2132, doi:10.1107/S0907444904019158 (2004).

51 Chen, V. B. MolProbity: all-atom structure validation for macromolecular crystallography. Acta Crystallogr. D 66, 12–21 (2010).

52 Smart, O. S., Neduvelil, J. G., Wang, X., Wallace, B. A. & Sansom, M. S. HOLE: a program for the analysis of the pore dimensions of ion channel structural models. J Mol Graph 14, 354–360, 376 (1996).

53 Harder, E. et al. OPLS3: A Force Field Providing Broad Coverage of Drug-like Small Molecules and Proteins. J Chem Theory Comput 12, 281–296, doi:10.1021/acs.jctc.5b00864 (2016).

54 Jo, S., Kim, T., Iyer, V. G. & Im, W. CHARMM-GUI: a web-based graphical user interface for CHARMM. J Comput Chem 29, 1859–1865, doi:10.1002/jcc.20945 (2008).

55 MacKerell, A. D. et al. All-atom empirical potential for molecular modeling and dynamics studies of proteins. J Phys Chem B 102, 3586–3616, doi:10.1021/jp973084f (1998).

56 Vanommeslaeghe, K. et al. CHARMM general force field: A force field for drug-like molecules compatible with the CHARMM all-atom additive biological force fields. J Comput Chem 31, 671–690, doi:10.1002/jcc.21367 (2010).

57 Berendsen, H. J. C., van der Spoel, D. & van Drunen, R. GROMACS: a message-passing parallel molecular dynamics implementation. Comput Phys Commun 91, 43–56 (1995).

58 Bussi, G., Donadio, D. & Parrinello, M. Canonical sampling through velocity rescaling. J Chem Phys 126, 14101 (2007).

59 Hoover, W. G. Canonical dynamics: Equilibrium phase-space distributions. Phys Rev A Gen Phys 31, 1695–1697 (1985).

60 Hess, B., Bekker, H., Berendsen, H. J. C. & Fraaije, J. G. E. M. LINCS: A linear constraint solver for molecular simulations. Journal of Computational Chemistry Volume 18, 1463–1472 (1997).

61 Humphrey, W., Dalke, A. & Schulten, K. VMD: visual molecular dynamics. J Mol Graph 14, 33-38, 27–38 (1996).

62 Van Der Spoel, D. et al. GROMACS: fast, flexible, and free. J Comput Chem 26, 1701–1718, doi:10.1002/jcc.20291 (2005).

63 Trendelkamp-Schroer, B., Wu, H., Paul, F. & Noe, F. Estimation and uncertainty of reversible Markov models. J Chem Phys 143, 174101, doi:10.1063/1.4934536 (2015).

